# Mapping antigenic evolution of influenza A virus using deep learning-based prediction of hemagglutination inhibition titers

**DOI:** 10.1101/2025.08.22.670987

**Authors:** Bingyi Yang, Yifan Yin, Lin Wang, Tim K. Tsang, Nicholas C. Wu, Henrik Salje

**Author notes:** Correspondence to: Bingyi Yang. These authors contributed equally (Bingyi Yang, Yifan Yin, Lin Wang).

## Abstract

Seasonal influenza remains a significant public health challenge through unpredictable antigenic drift, where accumulated mutations enable immune evasion and necessitate vaccine updates. Comprehensive antigenic characterization has been hindered by the absence of real-world A(H1N1)pdm antigenic mapping and post-2012 A(H3N2) maps due to insufficient pair-wise hemagglutination inhibition (HAI) titrations. Here, we present an end-to-end transformer model that predicts HAI titers directly from viral genetic sequences with error under two-fold, comparable to experimental variability. This approach enables rapid and high-throughput augmentation of HAI titrations across viral isolates, allowing for constructing large-scale antigenic maps. Several novel evolutionary patterns emerged from our analysis. Our antigenic mapping identified three A(H3N2) clusters between 2012-2022 with transitions occurring approximately every 6 years. Notably, genetically diverse co-circulating subclades 3C.2a and 3C.3a (2015-2020) formed a single antigenic cluster. A(H1N1)pdm formed three less temporally distinct clusters, with a novel cluster emerging post-COVID-19. Using interpretable analysis, we identified mutation sites linked to antigenic cluster transitions that align with previous laboratory findings. Key A(H3N2) mutations primarily occurred within major antigenic epitopes, while A(H1N1) showed fewer key mutations, some outside recognized antigenic regions. Our deep learning approach accelerates antigenic characterization in surveillance at global scale and can be transferred to other variable pathogens, providing an actionable bridge between genomic sequencing and vaccine strain selection.

## Introduction

Influenza epidemics impose significant public health burdens globally ^1^. Currently circulating influenza A viruses in humans consist of two major subtypes: A(H3N2) and A(H1N1)pdm. A(H3N2) emerged during the 1968 pandemic and has been circulating for nearly six decades, while A(H1N1)pdm replaced previous seasonal A(H1N1) strains after the 2009 pandemic. Both subtypes undergo continuous genetic and antigenic evolution, leading to cross-reactivity between previously encountered and contemporarily circulating strains ^2–6^. This cross-reactivity shapes pre-existing immunity and influences the protective effects of prior immune responses, necessitating precise monitoring of antigenic changes and regular updates to influenza vaccines to maintain effectiveness ^2,5–7^.

Antigenic mapping is an essential tool that quantifies viral evolution by projecting high-dimensional cross-reactivity data into two or three dimensions, guiding vaccine strain selection ^6^. For A(H3N2), the most comprehensive antigenic maps only include viruses isolated through 2002 (Figure 1D), with an update in 2014 ^4,6^. This represents a significant knowledge gap, as A(H3N2) genetic evolution changed markedly after 2013, characterized by the co-circulation and competition of two major clades (3C.2 and 3C.3) for almost a decade ^8^, in contrast to the mostly single-clade dominance observed from 1968 to 2012 (Figure 1C,E,F) ^3,4^. For A(H1N1)pdm, existing antigenic maps rely primarily on early 2009 pandemic strains and laboratory-generated mutants rather than circulating isolates ^5^, leaving a significant gap in our understanding of A(H1N1)pdm antigenic evolution since its emergence.

**Figure 1.**
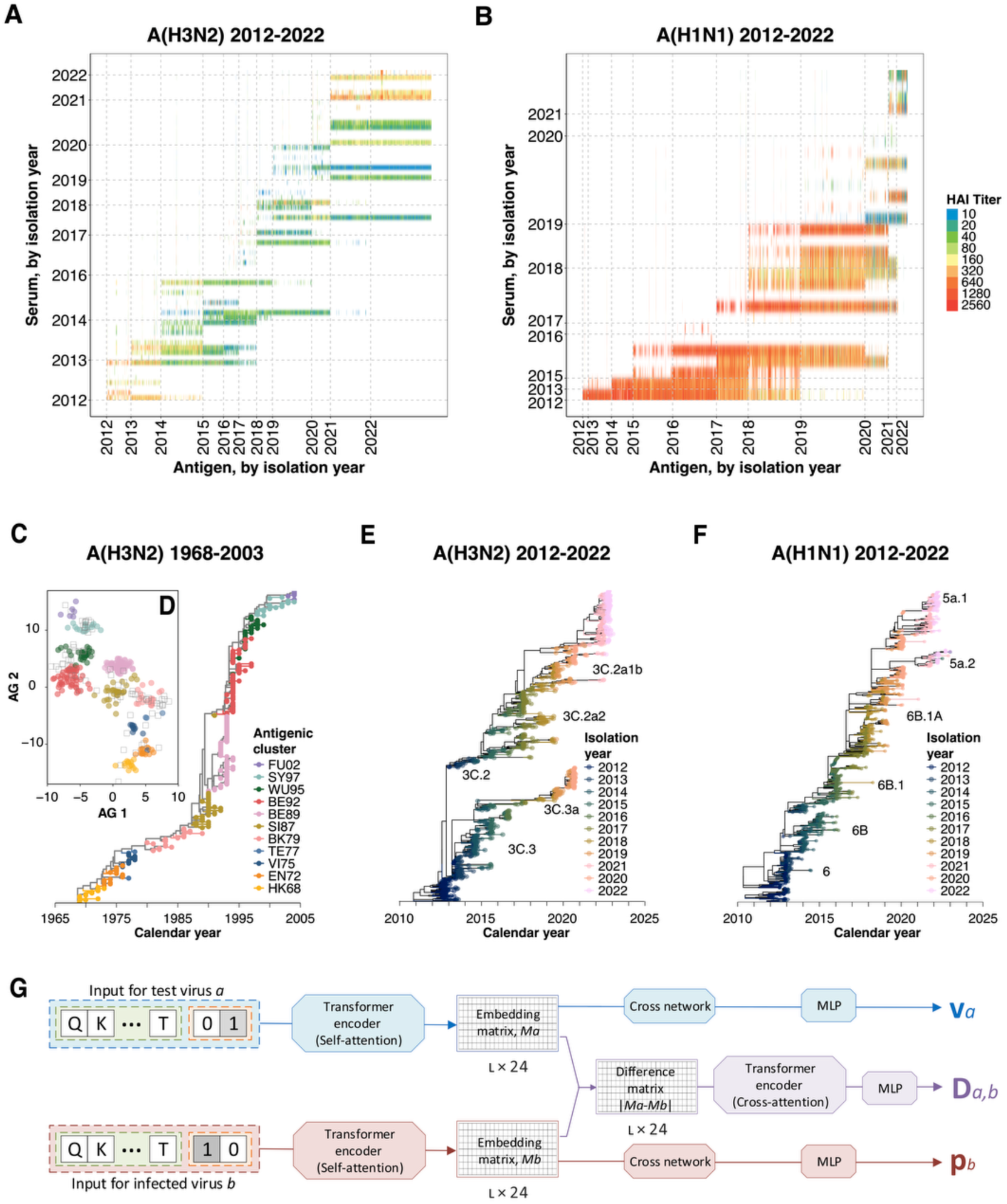
Predicting antigenic change from HA1 sequences with a deep-learning framework. (**A**) Cross-reactivity measured by hemagglutination inhibition (HAI) titers for A(H3N2) isolated between 2012-2022. In total 64,301 titers from 112 reference viruses and 4,154 test viruses were included. (**B**) Cross-reactivity measured by HAI titers for A(H1N1)pdm isolated between 2012-2022. In total 65,505 titers from 48 reference viruses and 4,803 test viruses were included. (**C**) Time-calibrated phylogenetic tree for A(H3N2) viruses isolated between 1968-2003, colored by antigenic clusters shown in panel **D**. (**D**) Antigenic map and clusters for A(H3N2) viruses isolated between 1968-2003, estimated by Smith *et al*. 2004 ^6^. (**E**) Time-calibrated maximum likelihood phylogenetic tree for A(H3N2) viruses isolated between 2012-2022. We randomly selected 880 strains with 80 for each year (same for panel F). (**F**) Time-calibrated maximum likelihood phylogenetic tree for A(H1N1)pdm viruses isolated between 2012-2022. (**G**) Schematic representation of the model used to predict HAI responses based on HA1 sequences from the test and reference viruses. The model predicts both log HAI titers and fold changes in titers, determined by the model-learned test virus avidity (*v_a_*), antiserum potency (*p_b_*), and genetic distance between the test and reference viruses (*D_a,b_*). For each virus-antiserum pair, sequence data (with length *L =* 329 for A(H3N2) and 327 for A(H1N1)pdm) and metadata are encoded into 24-unit vectors for both the test virus *a* and the reference virus *b*, which are processed through transformer encoders with self-attention to generate the encoded matrices *M_a_* and *M_b_*. A deep cross network and multi-layer perceptron (MLP) are then applied to these encoded matrices to derive *v_a_* and *p_b_*. Simultaneously, element-wise subtraction and absolute operations (|*M_a_* – *M_b_*|) are performed to derive the difference matrix, which is used alongside a transformer module with cross-attention mechanisms and MLP to calculate *D_a,b_*.

While genetic evolution can be tracked through sequence analysis, antigenic evolution specifically involves structural changes in epitopes that facilitate escape from antibody recognition and does not always mirror genetic changes ^9,10^. Koel *et al*. identified seven key amino acid positions in antigenic epitopes A and B that drove antigenic cluster transitions in A(H3N2) viruses from 1968 to 2002 ^11^. Although studies have linked specific mutations (e.g., positions 158-160) to altered antigenicity between subclades and potential vaccine mismatches after 2012 ^8,12^, the global impact of genetic co-circulation on A(H3N2)’s antigenic evolution remains unclear. For A(H1N1)pdm, while studies suggested that the Sb and Ca2 sites are immunodominant among the five recognized antigenic epitopes ^9,10^, the specific positions linked to antigenic cluster transitions in circulating strains remain largely incompletely identified ^5,11^.

A key challenge in characterizing antigenic evolution in post-2002 A(H3N2) and A(H1N1)pdm viruses is that antigenic mapping requires resource-intensive laboratory work, including ferret challenges and hemagglutination inhibition (HAI) assays. Meanwhile, the rapid expansion of genetic sequence data on platforms such as GISAID has created opportunities for *in silico* approaches ^13^. Researchers have developed various machine learning models that leverage genetic data to predict influenza antigenic evolutions. Early substitution models for A(H3N2) HAI titers showed promise but were constrained by their reliance on known substitutions ^14^. Alternative approaches using deep learning embeddings, physicochemical properties, or graph neural networks have also demonstrated potential but face challenges in computational efficiency, accuracy, flexibility, or data utilization ^15–18^. Notably, these predictive models have focused almost exclusively on historical A(H3N2) strains, with few comparable models for A(H1N1)pdm viruses or recent A(H3N2) strains and often do not account for factors like passage history. A more recent model using AdaBoost with amino acid (AA) substitution matrices (GIAG010101) for physicochemical encoding has shown improved performance for recent A(H3N2) data; however, its fixed training limits flexibility and rapid adaptation to newly isolated strains ^19^. This highlights the need for flexible, data-driven approaches that leverage expanding genetic data to improve surveillance and our understanding of contemporary antigenic evolution in both A(H3N2) and A(H1N1)pdm subtypes.

In this study, we aimed to characterize the antigenic evolution of influenza A(H3N2) and A(H1N1)pdm viruses from 2012 to 2022, using a novel end-to-end, transformer-based deep learning approach to overcome limitations in large-volume HAI titer data retrieved from the World Influenza Centre (WIC) at the Crick Institute ^20^. First, we developed a transformer model to predict HAI titers from sequences of the hemagglutinin (HA) protein and metadata from the WIC ^20^ and GISAID ^21^, achieving comparable or superior prediction accuracy and superior generalizability. Next, we applied this model to generate complete HAI titer matrices and construct high-resolution, subtype-specific antigenic maps, revealing distinct clustering patterns that were previously obscured by incomplete HAI data. Finally, we identified key AA positions that are associated with antigenic cluster transitions through interpretable analysis of our deep learning model, comparing these findings with known antigenic sites and validating them against previous laboratory results to enhance our understanding of influenza antigenic evolution.

## Results

### Sequence and titer data of A(H3N2) and A(H1N1)pdm viruses

We compiled HAI titers for A(H3N2) and A(H1N1)pdm viruses using the biannual reports of influenza surveillance and vaccine recommendations from the WIC from 2012 to 2023 ^19^. Titers measured using ferret antisera raised against reference viruses were tested against viruses. Most test viruses (81% for A(H3N2) and 77% for A(H1N1)pdm) were isolated in the same year as or within three years after the isolation of the corresponding reference virus (Figure 1A-B). Information including virus name, isolation year, synthetic viruses, and propagation method was extracted, with viruses classified as egg-propagated if ever propagated in eggs, otherwise as cell-propagated. HA1 segment nucleic acid (NA) and AA sequences for A(H3N2) and A(H1N1)pdm viruses were retrieved from the WIC reports and GISAID ^20,21^. All AA sequences were used as input for the deep learning models, while 880 randomly selected NA sequences per subtype (80 per year) were used to construct time-calibrated phylogenetic trees.

The dataset included 64,301 titers for A(H3N2) (112 reference viruses, 4,154 test viruses) and 65,505 titers for A(H1N1)pdm (48 reference viruses, 4,803 test viruses) (Figure 1A-B). Viruses isolated in close temporal proximity to the reference viruses exhibited higher titers than those isolated earlier. For example, for A(H3N2), viruses isolated in the same year had a median titer of 320 (interquartile range (IQR) 160– 640), compared to 80 (40–160) for those isolated four years later (Figure 1A). Additionally, titers were generally higher for A(H1N1)pdm (median 1280, IQR 640– 2560, same-year isolation) than for A(H3N2) (median 320, IQR 160–640). The phylogenetic tree revealed co-circulation of clade 3C.2a and 3C.3a since 2014 for A(H3N2) and an emerging clade 6B.1A.5a.2 for A(H1N1)pdm in 2021 (Figure 1E-F).

### Deep learning model

#### End-to-end model structure

We adapted the deep cross network (DCN) model using transformer encoder-processed AA sequences and other viral information to learn the relationship between protein sequences and HAI responses in a supervised manner. Our model predicts log-transformed HAI titers 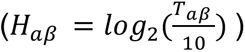, which measure antibodies against test virus *a* of antiserum *β* raised from a reference virus *b*. Following previous studies, we assumed that antigenic responses were determined by three components: virus avidity *v*_*a*_, antiserum potency *p*_*b*_, and genetic difference *D*_*ab*_ (Figure 1G) ^14^, with the model prediction expressed as:

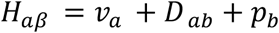

Instead of counting the AA differences that assumed each site are independent and equally important, we incorporated the entire HA1 protein sequences of both test and reference viruses as model inputs to derive *v*_*a*_, *p*_*b*_ and *D*_*ab*_, allowing the model to learn the relationship between antigenic response and predictors (Figure 1G). We also included information on passage history and synthetic virus for both viruses as model input (Methods).

Each virus (test virus *a* or reference virus *b*) was represented by *L* variables (*L* is 331 for A(H3N2) and 329 for A(H1N1)pdm), with each variable embedded into a 24-unit vector (Figure 1G). Using a transformer encoder with multi-head self-attention, we obtained a *L* × 24 embedding matrix (*M*_*a*_ or *M*_*b*_) that captures positional information and AA interactions. These matrices underwent training using DCN to calculate virus avidity *v*_*a*_ and antiserum potency *p*_*b*_ (Figure. 1G).

The difference between test and reference virus embedding matrices was processed through a transformer module with cross-attention to generate a difference matrix (|*M*_*a*_ − *M*_*b*_|), which was input into a 3-layer MLP (512 units per layer) to compute the genetic difference *D*_*ab*_ (Figure 1G). All embedding weights were initialized with a normal distribution (mean = 0, SD = 1).

As a sensitivity analysis, we also trained the model using fold-change between test viruses and homologous titers (detailed in Methods).

#### The model accurately predicts HAI responses

We assessed the performance of our model in predicting HAI titers for A(H3N2) and A(H1N1)pdm separately (Figure 2A-B). We used data from 2012–2020 for ten-fold cross-validation, and reserved 2021–2022 for forecasting assessment. Data used for cross-validation includes 47,659 titrations from 89 reference and 2,915 test viruses for A(H3N2), and 60,791 titrations from 38 reference and 4,490 test viruses for A(H1N1)pdm. For each subtype, we randomly split the virus–antiserum pairs into ten equal folds for cross-validation. Our model achieved robust predictive accuracy in ten-fold cross-validation, with mean absolute error (MAE) consistently below 1 log-unit (equivalent to a two-fold difference in HAI titers). In particular, 98% of predictions fall within a four-fold titer difference for both subtypes (Figure 2C, E).

**Figure 2.**
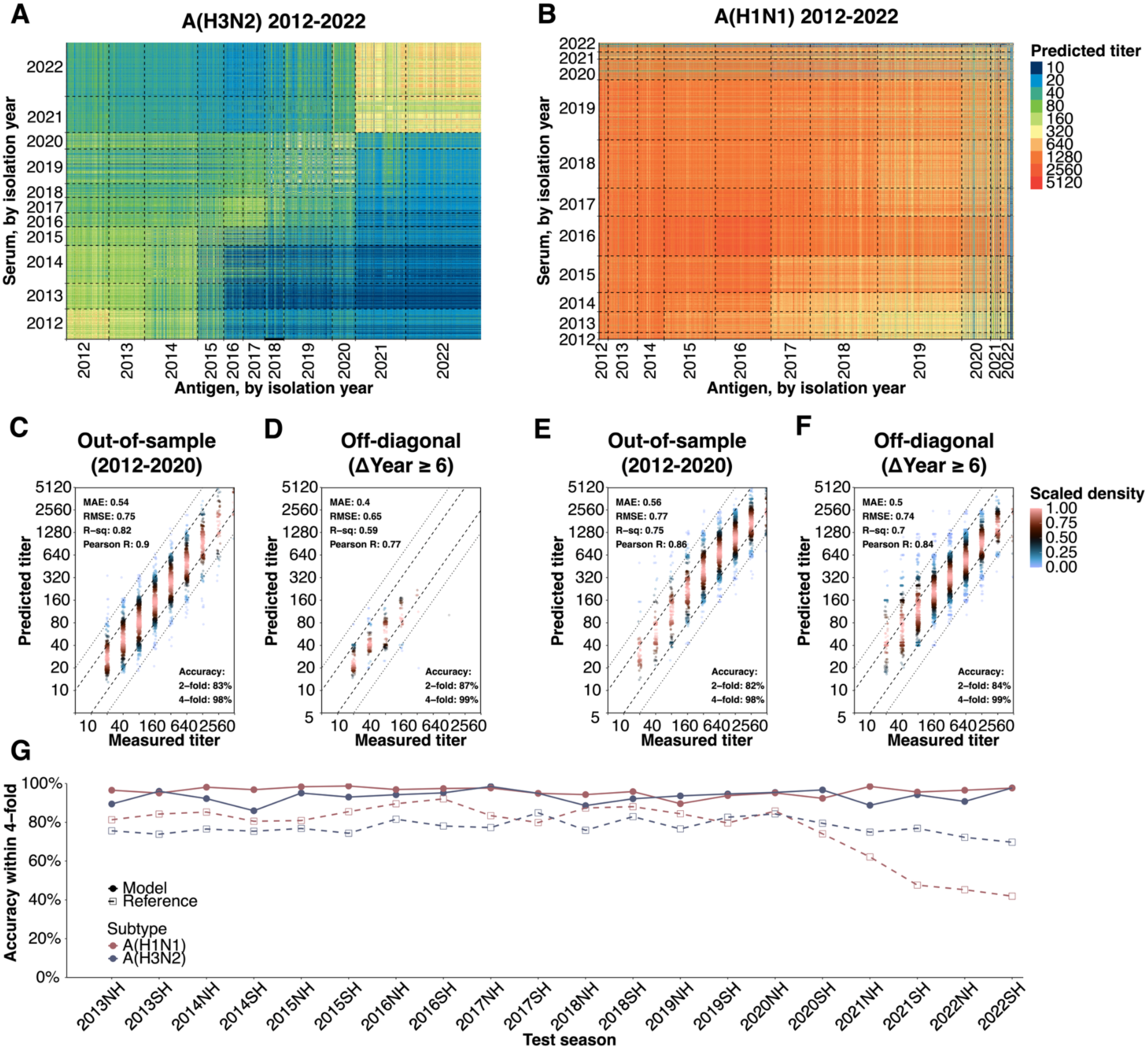
Model performance in predicting HAI titers. (**A**) Predicted HAI titers for A(H3N2) viruses isolated between 2012–2022, based on 64,301 titers from 112 reference viruses and 4,154 test viruses. Thick black lines in both panels represent diagonal and off-diagonal boundaries (same for **B**). (**B**) Predicted HAI titers for A(H1N1)pdm viruses isolated between 2012–2022, based on 65,505 titers from 48 reference viruses and 4,803 test viruses. (**C**) Ten-fold out-of-sample validation for A(H3N2) viruses isolated between 2012–2020. Dashed and dotted lines represent 2-fold and 4-fold prediction errors, respectively (same for **D**). (**D**) Off-diagonal HAI titer predictions for A(H3N2). (**E**) Ten-fold out-of-sample validation for A(H1N1)pdm viruses isolated between 2012–2020. (**F**) Off-diagonal HAI titer predictions for A(H1N1)pdm. (**G**) Prospective season-by-season predictions for A(H3N2) and A(H1N1)pdm, showing accuracy within 4-fold. Each point represents a model trained on data available prior to the given season, predicting titers for that season. Reference titers represent average titers measured in the corresponding season.

In the compiled surveillance dataset, most measurements lie on the diagonal, where virus–antiserum pairs were isolated ≤ 6 years apart. Off-diagonal titers (> 6 years) account for only 0.8 % in A(H3N2) and 23 % in A(H1N1)pdm (Figure. 1A, B). Our model achieved high accuracy for these off-diagonal pairs, with 99% of titers predicted with <4-fold difference for both subtypes (Figure 2D, F). This underscores the model’s ability to predict antigenic cross-reactivity between viruses with substantial temporal gaps. Using fold-of change as model outcome and additional metrics including root mean square error, R-squared, and Spearman correlation, showed consistent robustness and accuracy of our model performance (Figures 2C, E and S1-S2).

Our model demonstrated accurate nowcasting performance in season-to-season rolling prediction assessments. For each season, we trained the model using sequence and titer data from all preceding seasons ^19^. Our model achieved 86–98% accuracy for A(H3N2) and 90–99% for A(H1N1)pdm within a four-fold difference of measured HAI titers, outperforming those based on the previous season’s average antisera titers (Figure 2G). Notably, our model maintained high-level prediction accuracy despite the significant reduction in A(H1N1)pdm cross-reactivity from the 2020 south hemisphere season. Our model had consistent performance in predictive accuracy when assessing with other performance metrics and when using fold-change as the model output (Figures S3–S4).

Our model maintained a high-level predictive accuracy for viral isolates from up to two years ahead, achieving 85% accuracy for A(H3N2) and 96% for A(H1N1)pdm within a four-fold difference (Figure S1 C,F). Compared to the substitution-based and machine learning models ^14,19^, our approach provided improved (or comparable) performance in cross-validation and season-by-season nowcast assessments, which holds for both subtypes across all evaluation metrics (Figures S5–S6).

### Antigenic mapping using HAI predictions reveals clustering and evolutionary patterns

#### Complete titer matrix reveals distinct antigenic clustering patterns

We first constructed subtype-specific antigenic maps using measured HAI titers for reference and test viruses from 2012 to 2022 (“incomplete”; Figure 3A), with a small proportion of off-diagonal titers (Figure 1A-B). Two-dimensional antigenic maps generated using multidimensional scaling (MDS) ^6,22^ showed limited clustering patterns (Figure 3D,G and Table S1) despite strong correlations between map and measured titer distances (Spearman correlation: 0.93 for A(H3N2) and 0.95 for A(H1N1)pdm with p<0.01; Figure S7A,D). For example, the optimal five-cluster solution for the incomplete A(H3N2) matrix yielded only a modest silhouette score of 0.46, indicating weakly separated and overlapping clusters (Table S1).

**Figure 3.**
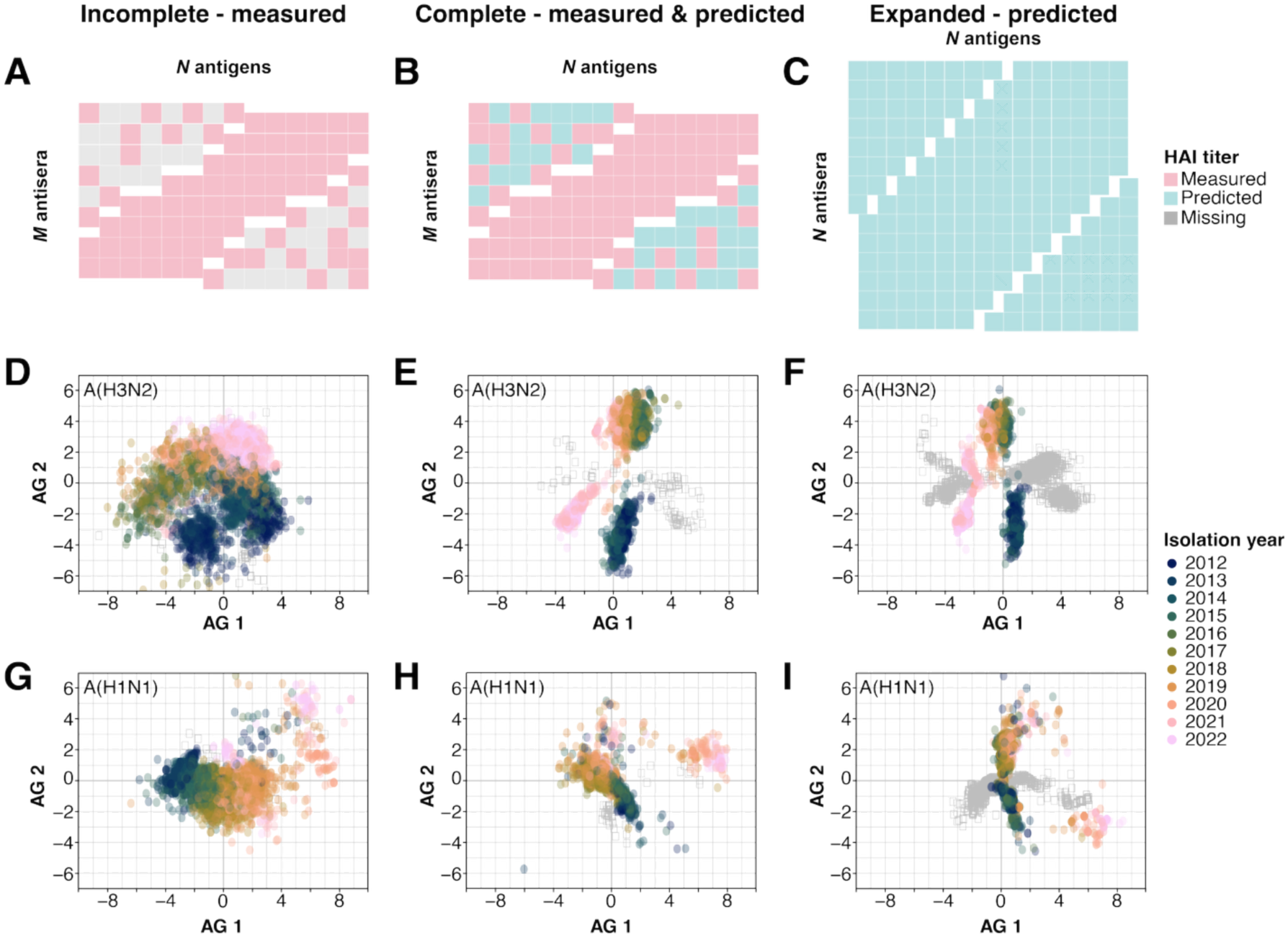
Antigenic maps for influenza A(H3N2) and A(H1N1)pdm viruses isolated between 2012 and 2022. (**A-C**) Conceptual illustration of HAI titer table compositions used for antigenic mapping. **(A)** An incomplete titer table representing measured HAI titers for *M* antisera against *N* viruses (*M<N*). Grey cells indicate unavailable data for specific virus-antiserum pairs. **(B)** A complete titer table combining measured titers with predicted values for the missing entries shown in panel A. **(C)** An expanded titer table displaying predicted HAI titers for all possible combinations of *N* antisera against *N* viruses, covering full coverage of antigen interactions. (**D-F**) Antigenic maps for A(H3N2). **(D)** Map fitted with the incomplete measured titer table (Figure 1A). **(E)** Map fitted with the completed titer table (combining measured and predicted titers). **(F)** Map fitted with the expanded predicted titer table, covering all virus-antiserum pairs (Figure 2A). (**G-I**) Antigenic maps for A(H1N1)pdm. **(G)** Map fitted with the incomplete measured titer table (Figure 1B). **(H)** Map fitted with the complete titer table (combining measured and predicted titers). **(I)** Map fitted with the expanded predicted titer table, covering all virus-antiserum pairs (Figure 2B).

When we generated antigenic maps by filling missing off-diagonal values in the cross-reactivity data with HAI titer predictions (“complete“; Figure 3B), the resulting maps displayed clear clustering patterns for both subtypes (Table S1) while maintaining significant correlations between map and titer distances (0.85 for A(H3N2) and 0.88 for A(H1N1)pdm, p<0.01; Figure S7B,E). Further expansion of the analysis by treating all test viruses as reference viruses (“expanded“; Figure 3C) produced similar clear clustering patterns with strong correlations between map and titer distances (0.92 for A(H3N2) and 0.92 for A(H1N1)pdm, p<0.01; Figure S7C,F). Specifically, the expanded dataset produced a consistent three-cluster solution for both A(H3N2) and A(H1N1)pdm, with higher silhouette scores (0.78 and 0.80) and supported by Calinski–Harabasz and Davies–Bouldin values that point to well-separated clusters (Table S1).

We observed similar discrepancies in antigenic maps generated using model-predicted A(H3N2) HAI titers for both incomplete versus complete matrices (Figure S8), indicating that filling missing data facilitates clearer clustering patterns. This mirrors the geographic analogy, where using only distances between nearby cities fails to recover true spatial relationships, but including all pair-wise distances accurately reconstructs them (Figure S9). Expanding the reference set of viruses or filling missing data substantially improved agreement between predicted antigenic distances from independent runs and data sets (Figures S10–S11).

We evaluated higher-dimensional antigenic maps. Although additional dimensions tend to reduce stress values, antigenic distance from 2D, 3D, and 4D maps were highly correlated (r > 0.95, p<0.01; Figure S12). For simplicity and clarity, we used 2D maps for all subsequent analyses.

#### Antigenic evolution of influenza A(H3N2)

Using antigenic maps derived from expanded titer matrices, we identified three distinct clusters among A(H3N2) viruses circulating between 2012 and 2022 (Figure 4A–C). Cluster 1 had a median isolation year of 2013 (range 2012–2015), cluster 2 a median of 2018 (range 2014–2022), and Cluster 3 comprised isolates collected from 2021 onward.

**Figure 4.**
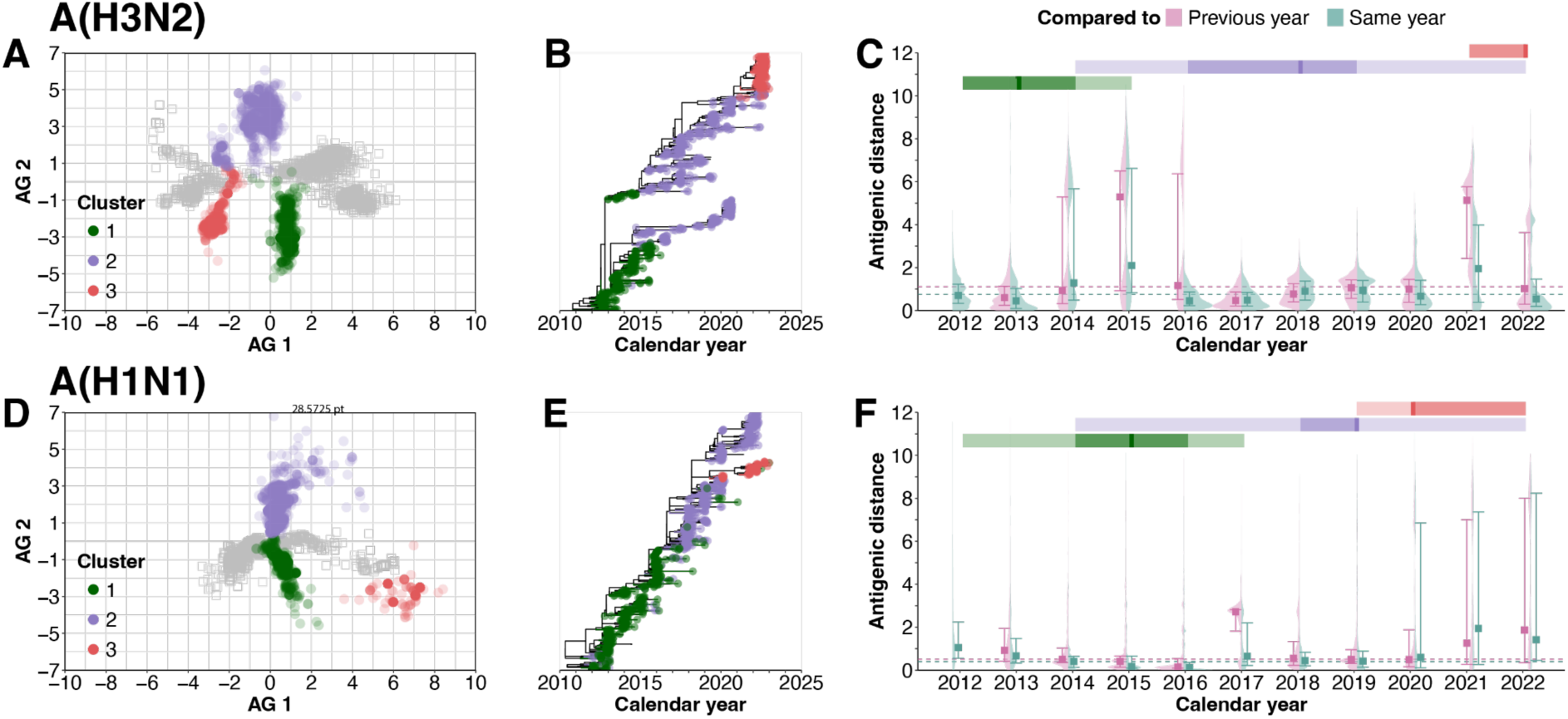
Antigenic evolution of influenza A(H3N2) and A(H1N1)pdm viruses, 2012–2022. (**A, D**) Antigenic maps derived from the expanded titer matrices (see Figure 3F, I) reveal discrete clusters for A(H3N2) (**A**) and A(H1N1)pdm (**D**). Clusters were identified using Gaussian Mixture Models (see Methods). (**B, E**) Time-scaled maximum-likelihood phylogenies (see Figure 1E,F), with branches colored by the clusters defined in panels A and D for A(H3N2) (**B**) and A(H1N1)pdm (**E**). (**C, F**) Pair-wise antigenic distances within each calendar year (green) and between each year and the preceding year (red) for A(H3N2) (**C**) and A(H1N1)pdm (**F**). Horizontal bars above each plot summarize the temporal span of every cluster; the thick line marks the median isolation date, the dark band the interquartile range, and the light band the 2.5th–97.5th percentiles.

We examined temporal movement in antigenic evolution by calculating pair-wise distances between strains isolated in the same year and comparing them to strains from the previous year. During the persistent period of antigenic clusters (typically near each cluster’s median year), viruses display close antigenic relationships. For instance, in 2013, strains exhibited median antigenic distances of 0.5 log-unit (IQR 0.1–1.0) to contemporaneous isolates and 0.6 log-units (IQR 0.3–1.1) to strains from the previous year (Figure 4C). In contrast, during cluster transition periods (2014– 2015 and 2021–2022), the distributions become distinctly bimodal. For example, among strains isolated in 2014, we observed a wider antigenic distance of 1.3 log-units (IQR 0.5–5.7) to other 2014 isolates, while maintaining closer relationships (median 0.6, IQR 0.2–1.0) to 2013 strains. This reflects the coexistence of antigenically distinct populations during these transition periods, with clear separation between intra-cluster and inter-cluster relationships (exemplified by the 2014 inter-cluster median of 5.9, IQR 5.4–6.5).

Despite genetic analyses identifying two subclades (3C.2a and 3C.3a) since 2014, our antigenic mapping grouped these subclades into a single cluster, distinct from isolates collected before 2013 and after 2021 (Figure 4A-B, S13A). Although cross-subclade HAI titers were generally lower than intra-subclade titers for these two subclades, median titers remained between 40 and 160 for contemporaneous strains, above the seropositivity threshold (Figure S14). Further, comparing antigenic distances revealed that divergence between 3C.2a and 3C.3a subclades was comparable to the typical antigenic difference between strains isolated one year apart, which is a rough threshold indicating an antigenic split for A(H3N2)-like virus (Figure S15) ^23^. Conversely, antigenic distances separating these two subclades from other clades exceeded this annual drift reference. Thus, our findings indicate that despite genetic divergence, antigenic profiles of 3C.2a and 3C.3a remain sufficiently similar to represent a single antigenic grouping, while earlier and recent isolates differ enough to form distinct clusters.

#### Antigenic evolution of influenza A(H1N1)pdm

For A(H1N1)pdm viruses from 2012 to 2022, our antigenic mapping also identified three distinct clusters: Cluster 1 centered around 2015 (range 2012–2017), Cluster 2 around 2019 (range 2014– 2022), and Cluster 3 emerging after 2019, predominantly comprising clade 5a.2 isolates (Figure 4D-F). Similar to A(H3N2), viruses collected near each cluster’s median year demonstrated stronger antigenic similarity to contemporaneous or recent strains (Figure 4F).

During transition periods (2016-2017 and 2020-2022), antigenic distance distributions exhibited bimodal patterns with peaks representing both intra-cluster and inter-cluster distances. However, A(H1N1)pdm clusters showed more extended transition periods with greater temporal overlap compared to A(H3N2). The transition between Clusters 1 and 2 in 2017 resulted in antigenic distances of approximately 3 log-units, while the transition from Clusters 2 to 3 produced more substantial antigenic divergence, with distances reaching up to 7 log-units (Figure 4F).

### Substitutions linked to antigenic cluster transitions

#### Interpretable approach to identify substitutions linked to antigenic cluster transitions

To identify AA substitutions associated with antigenic cluster transitions in influenza A viruses, we applied our deep learning model to representative viruses selected from WIC reports. These strains included vaccine candidates or dominant circulating viruses used in ferret serological challenge studies (Figure S16-17). For each antigenic cluster transition, including cluster 1 to 2 and 2 to 3 for A(H3N2), and cluster 1 to 2 for A(H1N1)pdm (based on data availability), viruses from the older cluster were used as reference (antisera), and those from the newer cluster as test viruses.

For each virus-antiserum pair, we applied Gradient-weighted Class Activation Mapping (Grad-CAM) to calculate saliency scores at each AA position. These scores quantified the contribution of individual positions to the predicted HAI titer by analyzing gradients and activations in the model’s cross-attention layer. For each antigenic transition, we averaged saliency scores across all virus-antiserum pairs at each variable position, defined as positions with observed substitutions. We then normalized these scores relative to the mean score across all variable positions. Positions with normalized scores greater than 1 were classified as key sites, reflecting their above-average contribution to antigenic cluster transitions.

To evaluate the biological relevance of these key sites, we examined their distribution across known major antigenic epitopes in the HA1 ^24^. For A(H3N2), we considered major antigenic epitopes A–E, and for A(H1N1)pdm, Sa, Sb, Ca1, Ca2, and Cb (Figure 5A-C; Figures S16-17). We then calculated the relative risk (RR) of observing key sites within each epitope compared to random expectation, identifying regions disproportionately involved in antigenic evolution (see Methods). For A(H3N2), we also calculated RR for the seven key sites (Figure 5A-B) identified by Koel et al. ^11^ for previous cluster transitions between 1968-2002.

**Figure 5.**
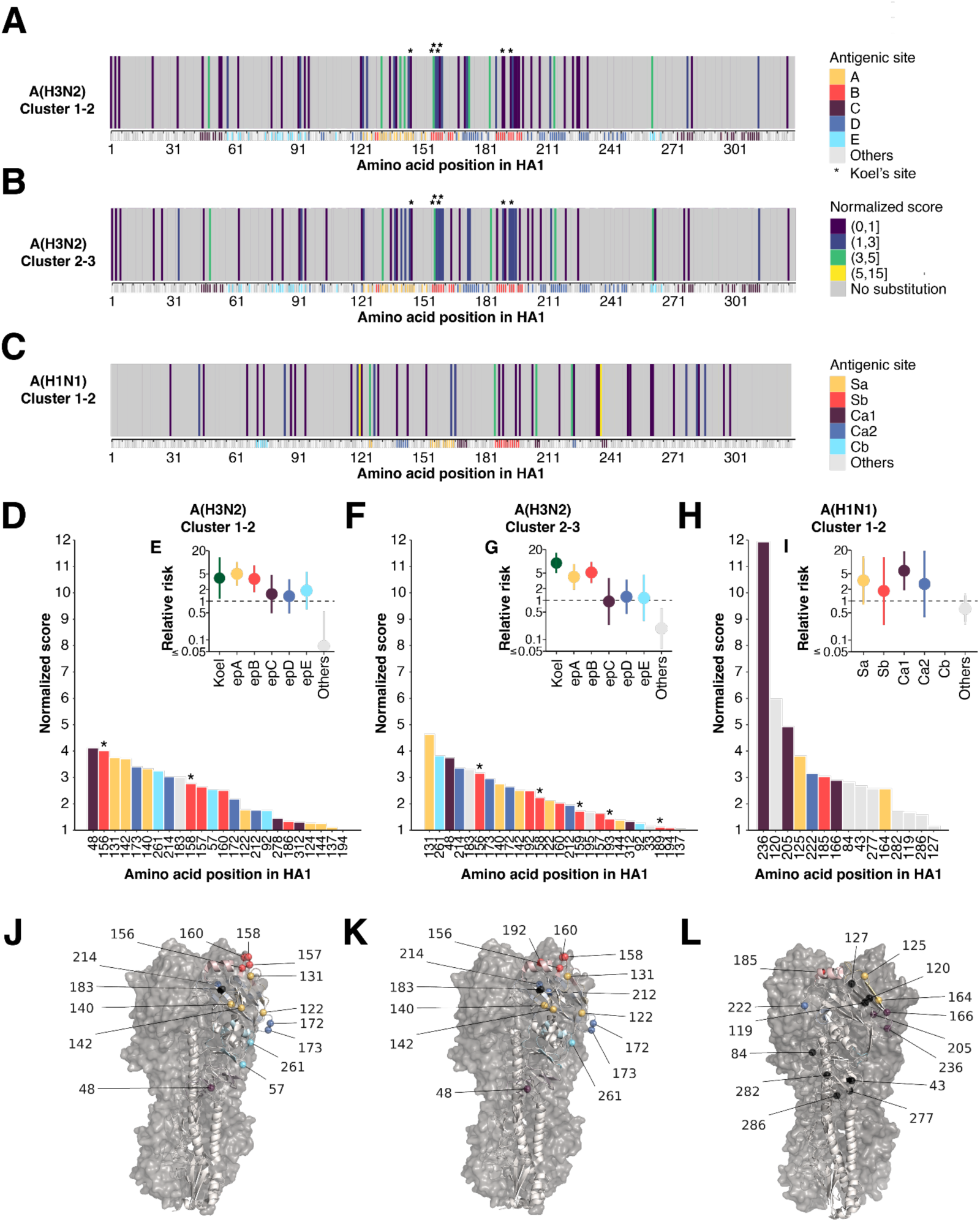
Model inferred key substitutions linking to antigenic cluster transitions for influenza A(H3N2) and A(H1N1)pdm. **(A-C).** Position-wise relative importance of embedding differences across the HA1 sequence, with the x-axis shaded by canonical antigenic epitopes. Panels show transitions A(H3N2) cluster 1 → 2 (**A**), A(H3N2) cluster 2 → 3 (**B**), and A(H1N1)pdm cluster 1 → 2 (**C**). Importance values are normalized so that 1 equals the global mean. (**D, F, H**)Sequence positions which normalized importance exceeds 1, highlighting positions most strongly implicated in each transition. (**E,G,I**) Relative risk of identifying key sites from each epitope compared to random selection (see method). (**J-L**) Top fifteen most important substitution sites in each antigenic cluster transition mapped onto the HA protein structure, colored based on their corresponding epitopes (black if falls outside the epitopes). A(H3N2) and A(H1N1)pdm viruses were displayed using the HA structure of A/Aichi/2/1968 (PDB: 2YPG) and A/California/04/2009 (PDB: 3UBE).

For validation, we trained an independent transformer model using historical A(H3N2) sequences and HAI titer from 1968-2003 ^6^ (MAE 0.77 for 10-fold validation). Grad-CAM analysis of these earlier antigenic transitions revealed key substitutions consistent with those reported by Koel et al. and other experimentally validated sites (Figure S18) ^6,11^. As a negative control, positions with normalized scores lower than 0.01 rarely overlapped with known critical sites revealed by Koel *et al* or other laboratory experiments (Table S2), supporting the specificity of our method in identifying candidate sites for further laboratory validation.

#### Key substitutions linked to A(H3N2) cluster transitions

We identified 67 and 65 variable AA sites during the first (cluster 1 to 2) and second (cluster 2 to 3) antigenic cluster transitions, respectively (Figure 5A, B and Figure S16). Among these, 30 sites (24 in the first and 26 in the second cluster transition; 20 shared by both) had normalized saliency scores >1, implying substantial association with antigenic movement (Table S3). Ninety percent (27/30) were within known antigenic epitopes A–E, with 9 of the top 10 most important sites consistent across both transitions (Figure 5J, K).

Key sites identified during 2012-2022 transitions were predominantly within antigenic epitopes A (RR: 5.1, 95% CI 2.5-10.2 and 4.0, 95% CI 1.9-8.5 for the first and second transitions) and B (RR: 3.7 95% CI 1.7-8.2 and 5.2, 95% CI 2.8-9.7) (Figures 5E,G,I), consistent with laboratory findings ^11^. Additionally, positions adjacent to the receptor-binding site (156, 158, 159, 189, 193) identified historically by Koel *et al.* ^11^ showed strong signals during recent transitions (RR: 3.9, 95% CI 1.1-13.4; Figure 5E), suggesting the recent antigenic evolution of A(H3N2) virus follows previously described molecular mechanism.

#### Key substitutions linked to A(H1N1)pdm cluster transitions

We mainly analyzed the transition from the first (2012–2017) to the second cluster (2018–2022), as reference viruses were absent for the most recent A(H1N1)pdm cluster in 2022. We detected 61 positions with AA substitutions and identified 15 key sites with normalized saliency scores >1 (Figure 5C, Figure S17 and Table S4). Seven (47%) located in known antigenic sites (Sa, Sb, Ca1, Ca2, and Cb), including three in Ca1, which were 6.0-fold (95% CI, 2.0-17.6) more likely to observe key sites than expected by chance (Figure 5H-I).

Among the eight sites outside canonical antigenic regions, three (127, 277, 286) are potential N-glycosylation sequons, with change in 127 could affect accessing antigenic sites ^25,26^. Positions 119–120 sit in an α-helix adjacent to the receptor-binding site and have been linked to altered monoclonal-antibody recognition ^27^. Positions 43 and 282 fall within the head domain with no prior functional reports, while position 84 consistently separates swine and human A(H1N1) viruses from the 2009 pandemic lineage ^25,27^.

## Discussion

We have demonstrated the utility of our transformer-based learning model converting HA1 sequence data into antigenic insight. Across more than 100,000 serum-antigen pairs the model predicted HAI titers with an average error within two-fold dilution, outperforming existing substitution-based and machine-learning baselines. Filling the experimental matrix with these *in silico* titers exposed three antigenic clusters in both A(H3N2) and A(H1N1)pdm. For A(H3N2), the analysis unified the genetically divergent 3C.2a and 3C.3a lineages into a single antigenic space, while for A(H1N1)pdm it revealed a post-COVID-19 pandemic cluster dominated by clade 5a.2. Grad-CAM interpretation recovered positions in major epitopes A and B linking to A(H3N2) cluster transitions and identified a smaller set of substitutions positioned partly outside recognised antigenic sites that were associated with recent A(H1N1)pdm drift. Together, our findings demonstrate that sequence-to-titer modelling can deliver real-time, pathogen-wide antigenic mapping and accelerate vaccine-strain selection for influenza and other rapidly evolving viruses.

The transformer model predicts titers directly from virus–antiserum sequences, passage history and synthetic information, without predefined antigenic features or extensive prior knowledge. The model dynamically learned antigenic differences, virus avidity and antisera potency for individual viruses, eliminating the need for predefined properties. By incorporating both protein position and specific substitution information, it holds the potential for capturing non-linear interactions for position-substitution and enhanced explanatory power that traditional substitution matrices miss ^14,19^. This flexibility and high predictive capability allow the model being used for timely applications, particularly informing the biannual vaccine recommendations, complementing the few-selected ferret antisera. Notably, the model successfully captured recent antigenic changes in A(H1N1)pdm since 2020 ^28^, underscoring its potential for real-time nowcasting of HAI titers and supporting vaccine recommendations by predicting titers for newly isolated viruses.

By augmenting incomplete diagonal HAI data with model-predicted off-diagonal titers, we produced a complete pair-wise matrix that is critical to further resolved previously obscured antigenic clusters. Our results, in line with previous work ^29^, show that even below-threshold titer information stabilizes cluster inference, much as incorporating all pair-wise distances restores geometric fidelity in a city-map reconstruction (Figure S9). These results suggest that sequence-based machine-learning models trained directly on antigenic distances should benchmark their outputs against maps built from fully covered titer matrices to avoid systematic bias ^15–17^.

Antigenic mapping revealed three clusters in both A(H3N2) and A(H1N1)pdm, yet their drift trajectories differed. For A(H3N2), the dominant cluster changed roughly every 3–4 years, consistent with historical patterns ^6^. The co-circulating 3C.2a and 3C.3a lineages formed a single antigenic group, supporting reports that their cross-reactivity differs more in magnitude than in breadth ^7,30^ and demonstrating that genetic divergence does not always translate into antigenic separation ^31^. A(H1N1)pdm showed a prolonged overlap between its first two clusters, reflecting slower mutation and broader cross-reactivity ^3,7,30^, followed by an abrupt jump to clade 5a.2 that has been linked to reduced vaccine effectiveness and a subsequent strain update ^28,32^. For both subtypes, we found stage-specific patterns for pair-wise antigenic distances among viruses isolated in the same year. During periods of cluster stability, the distribution was unimodal and centred on small distances, whereas during transition years it became bimodal, retaining the small-distance peak and adding a second peak at larger distances. These movement signatures, combined with the augmented HAI matrices, could improve real-time antigenic characterization, though achieving reliable forecasts will require more representative viruses and methods that account for the influence of future isolates on current antigenic positions.

Our analysis suggests that antigenic cluster transitions in A(H3N2) remain mostly linked to substitutions at the surface epitopes A and B, aligning with laboratory evidence of historical cluster transitions ^11^. In contrast, recent A(H1N1)pdm viruses show fewer key AA substitutions, many outside the known antigenic epitopes, including positions near the receptor-binding site and at the head-to-stem junction ^26,27^. These findings are consistent with the report that A(H1N1)pdm antigenic phenotype change depends on mutations that work in concert with receptor-binding sites rather than substitution in receptor-binding sites alone ^33^. Several A(H1N1)pdm key substitutions appeared in only one strain yet ranked highly in Grad-CAM analysis, and a full-sequence scan indicated that even conserved, structurally stabilising positions could affect antigenicity or fitness if altered. Because position effects are shaped by the local fitness landscape ^34^, we view these sites as provisional candidates rather than universal markers of drift. Our computational pipeline narrows the search space for critical mutations, but confirmatory reverse-genetics and binding assays are still required to establish their functional significance.

Our study has several caveats. First, while predicted off-diagonal HAI titers closely match available measurements and align with reports of minimal cross-reactivity among A(H3N2) strains more than six years apart ^7^, most of these predictions still await laboratory confirmation. Second, cluster assignments for recent viruses are provisional because right censoring means future isolates might shift their antigenic positions, so additional methods are needed to address this uncertainty. Third, Grad-CAM saliency reflects correlation rather than causation in antigenic cluster transitions, so high-scoring positions, especially those found in single A(H1N1)pdm isolates, need validation through reverse-genetics and binding assays. Fourth, because the model was trained only on HA1 sequences and HAI titers, interactions involving other genomic segments or entire viral genomes remain to be investigated. Finally, we could not analyse mutations linked to the latest A(H1N1)pdm cluster because post-2022 antigenic data were not yet compiled when this study was completed.

We demonstrated that a transformer model predicts influenza A HAI titers from HA1 sequences with two-fold accuracy, allowing us to impute missing off-diagonal values and assemble complete titration matrices. These matrices yielded distinctly clustered antigenic maps that resolved subtype-specific drift trajectories, and, through interpretable approaches, highlighted substitutions potentially linked to those shifts. This sequence-to-antigenic pipeline can complement laboratory surveillance and vaccine recommendation for influenza and, with further validation, may be adaptable to other antigenically variable viruses.

## Methods

### Hemagglutination-inhibition (HAI) data

We compiled data on A(H3N2) and A(H1N1)pdm that were available from the WHO Collaborating Centre for Reference and Research on Influenza at the Francis Crick Institute ^20^ published between 2012 and 2022. These titers were measured using ferret antisera raised against specific influenza A viruses. The antibody levels of the antisera (reference virus) were tested against various viruses. We extracted information on the reference virus and viruses, including virus name, year of first isolation, and propagation method, when available. Viruses were categorized as egg-propagated if they were ever propagated in eggs, even if also propagated in cells. Viruses propagated only in cells were classified as cell propagated. Isolation year was inferred from the virus name, and synthetic viruses were assigned the isolation year of their source domain virus. All included studies used the HAI test to measure antibodies. For A(H3N2), guinea pig red blood cells (RBCs) were used in the Crick reports ^20^. No RBC information was reported for the HAI tests used for A(H1N1)pdm.

We included 64,301 titers (39,245 unique virus-antiserum pairs) for A(H3N2) from 112 reference viruses and 4,154 test viruses isolated between 2012 and 2022. Of these titers, 30,005 (46.7%) were from cell-propagated viruses, 23,218 (36.1%) from test cell-propagated viruses with reference viruses in eggs, 5,468 (8.5%) from egg-propagated viruses, and 5,610 (8.7%) from test egg-propagated viruses with reference viruses in cells. 583 (0.9%) of pairs involved synthetic reference and 791 (1.2%) involved synthetic viruses.

For A(H1N1)pdm, we included 65,505 titers (45,289 unique virus-antiserum pairs) from 48 reference viruses and 4,803 test viruses isolated between 2012 and 2022. Of these titers, 23,264 (35.5%) were from cell-propagated viruses, 29,709 (45.4%) from test cell-propagated viruses with reference viruses in eggs, 7,230 (11%) from egg-propagated viruses, and 5,302 (8.1%) from test egg-propagated viruses with reference viruses in cells. 481 (0.7%) of pairs involved synthetic reference and 173 (0.3%) involved synthetic viruses.

### Sequence data

We retrieved HA1 nucleic acid (NA) sequences for A(H3N2) and A(H1N1)pdm viruses from GISAID (ref) and amino acid (AA) sequences from Crick’s biannual reports. For viruses without reported AA sequences in Crick’s reports, they were derived by translating the corresponding nucleic acid sequences from GISAID. Sequences with missing or ambiguous nucleotides that prevented accurate AA determination were excluded.

For A(H3N2), we obtained 4,172 complete AA sequences and corresponding nucleic acid sequences. For A(H1N1)pdm, we obtained 4,803 complete AA sequences and corresponding nucleic acid sequences. These AA sequences were used as input for the deep learning model and a subset of nucleic acid sequences were used to construct the phylogenetic tree.

Additionally, we retrieved HA1 segment nucleic acid sequences for 272 of 273 A(H3N2) viruses (isolated between 1968 and 2023) reported in Smith et al. (2004) from GISAID. All nucleic acid sequences were aligned by subtype using MAFFT ^35^.

### Time-calibrated maximum likelihood phylogenies of influenza A isolates

To construct phylogenetic trees, we subsampled 880 viruses with available nucleic acid sequences for each subtype, randomly selecting 80 per isolation year from 2012 to 2022. Maximum likelihood phylogenies were generated using IQ-TREE v2.3.6 ^36^ for the aligned HA1 segment sequences, applying the best-fitting nucleotide substitution model identified by ModelFinder ^36,37^. Time-calibrated trees were inferred using TreeTime v0.9.2 ^38^, based on deviations in root-to-tip regression of genetic distance against sampling dates. Tree tips were coloured by virus isolation year and antigenic clusters identified from antigenic maps using the complete HAI titer table, respectively.

We also constructed a phylogenetic tree for A(H3N2) viruses between 1968 and 2023 reported in Smith *et al*. using the same method, with tree tips coloured by antigenic clusters identified in the previous studies ^6,38^.

### Deep learning model

To provide numeric input for HAI titer and fold change of HAI titer for A(H3N2) and A(H1N1)pdm prediction model, we converted AA in each position to an integer using our predefined dictionary (K -> 0, C -> 1, …, S -> 18, D -> 19). Accordingly, metadata including passage history and synthetic virus were appended to the sequence data, in which passage history was labelled to 1 if the virus had ever been propagated in eggs, and synthetic virus was labelled to 1. For instance, if a cell-propagated synthetic virus has HA1 sequence KKCSSD, the input of the virus is [0, 0, 1, 18, 18, 19, 0, 1].

The model embedded each AA of HA1 sequences, passage history and synthetic virus into a 24-unit vector. A transformer encoder with 8 heads self-attention was applied to this process to obtain positional information and AA interactions. Using these embeddings of test and reference virus, virus avidity *v*_*a*_ and antiserum potency *p*_*b*_ were computed through two different DCNs, and the difference matrix was computed by taking the absolute value of the difference between those embeddings. The genetic difference *D*_*ab*_ was computed through another transformer module with a cross-attention mechanism to the difference matrix. The final output was the sum of *v*_*a*_, *p*_*b*_ and *D*_*ab*_. The embeddings of AA, passage history and synthetic virus were initialized with a normal distribution (mean = 0, SD = 1). For the hyperparameters of both transformer modules in the model, we used sinusoidal embedding as the positional embedding and 96 as the dimensionality of feed-forward inner-layer. For the DCN, the number of cross layers was 1 for A(H3N2) and 3 for A(H1N1)pdm. For the hyperparameters of MLP in our model, we chose 3 as the number of hidden layers and 512 as the number of nodes in each hidden layer. All the activation functions in our model were ReLU functions. The model was trained using Adam optimizer and L1 loss. The random seed was set to 100, batch size to 16 and number of epochs to 40, while other hyperparameters of Adam optimizer were default. Warm-up was also used in our model, in which the learning rate increased linearly from 0.00001 to 0.0005 in the first 10 epochs and decayed exponentially (gamma = 0.9) in the remaining 30 epochs. Training was conducted using PyTorch 2.4 on an NVIDIA RTX A2000 12GB GPU.

### Antigenic map and clusters

To construct the antigenic map, we utilized multidimensional scaling (MDS) analysis of HAI titers, using the “Racmacs” package in R ^39^. The analysis incorporated both experimentally measured HAI titers from Crick’s reports (Figure 1A-B) and predicted titers generated by our deep learning model (Figure 2A-B). The inclusion of predicted titers specifically addressed missing pairs in the measured dataset, particularly for off-diagonal virus-antiserum pairs with temporal separation greater than six years, which are often absent in experimental measurements.

Antigenic clusters of the constructed map were identified using Gaussian Mixture Models (GMM), which uses a probabilistic framework to capture clusters with elliptical shapes and varying sizes. The number of clusters was determined by several metrics, including silhouette score (measuring cohesion and separation), Calinski-Harabasz index (measuring between-cluster dispersion), and Davies-Bouldin index (measuring cluster separation). We selected 3 clusters for A(H3N2) as all metrics indicated it provided the best performance (Table S1). For A(H1N1)pdm, no single cluster number was optimal (with 2 clusters yielding the best silhouette and 4 clusters the lowest Davies-Bouldin index), so clustering was plotted for all candidate cluster numbers (Table S1). The analysis was performed using the “mclust” package in R ^40^.

#### World city map analogy analysis

To validate our method, we conducted an analogy experiment using real-world geographic data. We randomly selected ten cities from each of five continents (Africa, Asia, Europe, North America and South America), for a total of 50 cities. We computed pair-wise Euclidean distances, *d*(*city*_1_, *city*_2_) using latitude and longitude, and derived a complete distance table. To mimic sparse serological data, we retained only the shortest 20% of the distances, yielding an incomplete distance matrix. Both the complete and incomplete distance tables were converted into “titer” tables using the formula:

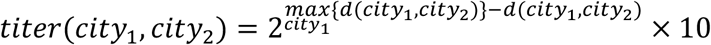

Where 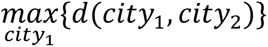 is the maximum distance from *city*_2_ to any city in the complete table. MDS was then applied to both titer tables using the “*Racmacs*” package in R ^39^, with the resulting maps subjected to translation and rotation for alignment. The antigenic map derived from the complete table closely resembled the original geographic distribution (Figure S9B-C). In contrast, the map generated from the incomplete table exhibited less distinct clustering, indicating reduced stability and clarity (Figure S9D-E).

### Model explanation analysis to identify key substitutions linked with antigenic cluster transitions

#### Virus-antiserum pair selection

To investigate antigenic cluster transitions, we selected representative viruses from two adjacent antigenic clusters. Viruses from the older cluster served as the reference (i.e., antisera), while those from the newer cluster were designated as the test viruses. These strains included vaccine candidates or dominant circulating viruses used in ferret serological challenge studies, including 76 for A(H3N2) and 30 for A(H1N1)pdm (Figure S16-17).

#### Position-wise saliency score for each virus-antiserum pair

For each representative virus-antiserum pair, we predicted the log-transformed HAI titer *H*_*aβ*_ using our deep learning model trained with all available data from 2012 to 2022 (Figure 1). We then performed backpropagation to compute gradients and activations of each model layer. Next, we applied Gradient-weighted Class Activation Mapping (Grad-CAM) ^41,42^ to the output of transformer module with cross-attention, which processes the absolute difference |*M*_*a*_ − *M*_*b*_| between the virus embedding matrices (Figure 1). This analysis quantifies the influence of the embedding difference at each sequence position *i* on the titer prediction. Specifically, let *K* denote the length of the embedding vector (with *K = 24*), er represented the output of the transformer module with cross-attention at position *i* as:

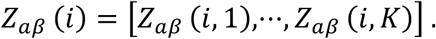

This vector *Z*_*aβ*_ (*i*) captures the positional contribution of the virus-antiserum embedding difference to the HAI titer prediction.

To calculate Grad-CAM using the predicted HAI titer *H*_*aβ*_ for each virus–antiserum pair, we first computed the gradient weight *ω*_*aβ*_ (*i*) at each AA position *i*. This weight indicates how changes in the embedding difference *Z*_*aβ*_ (*i*) affect the genetic difference *D*_*ab*_, using the following relation:

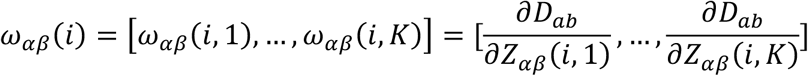

Next, we computed the weighted forward activation map *A*_*aβ*_ (*i*) by multiplying each gradient weight *ω*_*aβ*_ (*i*) with its corresponding embedding difference *Z*_*aβ*_ (*i*) and then summing across all dimensions *K*:

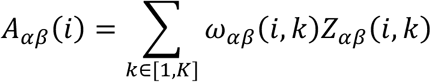

Finally, we computed a localization map Grad-CAM *S*_*aβ*_ (*i*) for each AA position *i* in the virus-antiserum pair, quantifying the impact of their difference on the predicted HAI titer:

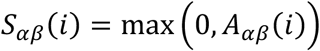

By doing such, it ensures that only positive contributions are considered. A higher saliency score indicates that the AA difference at position *i* has a stronger influence on the predicted titer, suggesting potential key sites involved in antigenic changes.

#### Averaging and normalizing saliency scores

To identify consistent patterns across multiple virus–antiserum pairs, we computed the average saliency score *S̅*(*i*) at each variable AA position *i* (defined as positions where substitutions occurred) across all relevant pairs involved in each antigenic cluster transition:

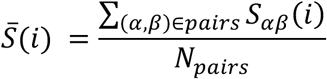

We then normalized these mean scores by the overall average saliency across all variable positions *V*, yielding a normalized measure:

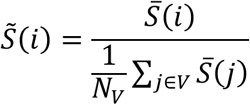

Positions with *S̃*(*i*) > 1 are considered key positions contributing above-average influence on antigenic transitions. Notably, we only considered positions that underwent substitutions during the transition, excluding conserved positions or those without observed changes during the study period. However, this exclusion does not imply that conserved positions are unimportant, as some may still exhibit high saliency scores.

#### Relative risk calculation

We calculated the relative risk (RR) of identifying a key position (i.e., with *S̃*(*i*) > 1) in each antigenic epitope *P* compared to a random selection among all key positions from the same antigenic epitope. We calculated the risk *R*_*P*_ of identifying a key position within antigenic epitope *P* using the number of key positions locating in antigenic epitope *P* (*n*_*P*_) and the number of all positions in the same antigenic epitope (*m*_*P*_):

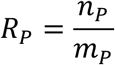

Similarly, we calculated the average risk of identifying a key position across the entire HA1 protein (*R*_*HA*1_):

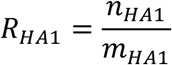

Finally, we derived the RR of identifying a key position for antigenic site *P* as follows:

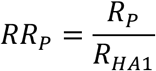

## Author contributions

B.Y.: Data compilation; deep-learning model conceptualization and execution; antigenic-cartography analysis; visualization; drafted the manuscript; funding acquisition. Y.Y.: Deep-learning model conceptualization and execution; Grad-CAM analysis; visualization. L.W.: Grad-CAM analysis; antigenic-cartography analysis; visualization; funding acquisition. T.K.T.: Data compilation; deep-learning model conceptualization; critical interpretation of results. N.C.W.: Critical interpretation of results. H.S.: Antigenic-cartography analysis; critical interpretation of results; funding acquisition.

## Acknowledgement

We thank the Worldwide Influenza Centre, Francis Crick Institute, UK for sharing antigenic data, especially Nicola Lewis and Ruth Harvey. We gratefully acknowledge all data contributors, i.e., the Authors and their Originating laboratories responsible for obtaining the specimens, and their Submitting laboratories for generating the genetic sequence and metadata and sharing via the GISAID Initiative, on which this research is based. This research was supported by the University of Hong Kong Seed Grant for Basic Sciences (Grant No. 2302101420) and Hong Kong Health and Medical Research Funds Research Fellowship Scheme (Grant No. 07210167) (to B.Y.). This research was supported by a subcontract from Johns Hopkins University with funds provided by Grant No. 75N93021C00045 from the National Institute of Allergy and Infectious Diseases, National Institutes of Health, Department of Health and Human Services (to H.S. and L.W.).

## Data and code availability

All data used in this study are publicly available from the Worldwide Influenza Centre, Francis Crick Institute, UK (https://www.crick.ac.uk/research/platforms-and-facilities/worldwide-influenza-centre), and GASID (https://gisaid.org). The source code for implementing the transformer model and Grad-CAM will be made openly available upon publication. The algorithm for multidimensional scaling (MDS) of antigenic maps was implemented using the R package “*Racmacs*,” which is publicly accessible at https://acorg.github.io/Racmacs/articles/intro-to-antigenic-cartography.html. Predicted HAI titers for A(H1N1) and A(H3N2) generated by our model are available on Hugging Face Spaces (https://huggingface.co/spaces/yyf031/Influenza_A_HAI_Titer_Prediction), where users can upload their own sequence pairs for prediction.

