## Supplementary materials for "Mapping antigenic evolution of influenza A virus using deep learning-based prediction of hemagglutination inhibition titers"

### Supplementary Tables

**Table S1.** Clustering evaluation metrics using partitioning around medoids (PAM) for varying numbers of clusters across measured, complete and expanded maps.

| Cluster number | Silhouette [a] | Calinski Harabasz score [b] | Davies Bouldin score [c] |
|---|---|---|---|
| **A(H3N2) incomplete (measured titers)** | | | |
| 2 | 0.43 | 3086 | 1.01 |
| 3 | 0.43 | 3779 | 0.80 |
| 4 | **0.46** | 4273 | 0.74 |
| 5 | **0.46** | 4672 | **0.69** |
| 6 | 0.44 | **4750** | 0.76 |
| 7 | 0.41 | 4407 | 0.80 |
| 8 | 0.40 | 4560 | 0.83 |
| **A(H3N2) complete (measured and predicted titers)** | | | |
| 2 | 0.74 | 17212 | 0.40 |
| 3 | **0.79** | **35744** | **0.28** |
| 4 | 0.61 | 28783 | 0.70 |
| 5 | 0.57 | 27890 | 0.72 |
| 6 | 0.51 | 26116 | 0.76 |
| 7 | 0.50 | 25308 | 0.77 |
| 8 | 0.52 | 27557 | 0.74 |
| **A(H3N2) expanded (predicted titers)** | | | |
| 2 | 0.69 | 12411 | 0.50 |
| 3 | **0.78** | **23046** | **0.28** |
| 4 | 0.66 | 24185 | 0.55 |
| 5 | 0.68 | 26127 | 0.53 |
| 6 | 0.68 | 23209 | 0.49 |
| 7 | 0.62 | 26145 | 0.53 |
| 8 | 0.62 | 27441 | 0.55 |
| **A(H1N1)pdm incomplete (measured titers)** | | | |
| 2 | **0.46** | 4229 | 0.83 |

| | | | |
|---|---|---|---|
| 3 | 0.38 | 3257 | 0.83 |
| 4 | 0.43 | **5353** | **0.74** |
| 5 | 0.39 | 5073 | 0.85 |
| 6 | 0.38 | 4937 | 0.83 |
| 7 | 0.36 | 4662 | 0.86 |
| 8 | 0.37 | 4430 | 0.86 |
| **A(H1N1)pdm complete (measured and predicted titers)** | | | |
| 2 | 0.53 | 3301 | 0.77 |
| 3 | **0.59** | **9063** | 0.49 |
| 4 | 0.55 | 8858 | **0.47** |
| 5 | 0.47 | 8529 | 0.66 |
| 6 | 0.49 | 8198 | 0.72 |
| 7 | 0.44 | 7385 | 0.75 |
| 8 | 0.43 | 6694 | 0.79 |
| **A(H1N1)pdm expanded (predicted titers)** | | | |
| 2 | 0.74 | 7925 | 0.44 |
| 3 | **0.80** | **19863** | **0.24** |
| 4 | 0.59 | 17749 | 0.57 |
| 5 | 0.62 | 17919 | 0.58 |
| 6 | 0.50 | 16420 | 0.72 |
| 7 | 0.52 | 15287 | 0.68 |
| 8 | 0.55 | 15422 | 0.66 |

a The silhouette score reflects how well-separated the clusters are, with values closer to 1 indicate better separation.

b The Calinski Harabasz index measures cluster dispersion, with higher values suggesting well-defined clusters.

c The Davies Bouldin score assesses cluster similarity and cluster separation, with lower values indicating better clustering performance.

**Table S2.** Model inferred sequence position with normalized scores less than 0.01 in historical antigenic cluster transitions, 1968-2002.

| Cluster transition | Antigenic epitopes | | | | | Other sites |
|---|---|---|---|---|---|---|
| | A | B | C | D | E | |
| HK68 to EN72 | 133,143 | 160 | 273,278 | 171,244 | 83 | 10,31, 34,139,264, 292 |
| EN72 to VI75 | 140 | 160 | 278(▼) | -- | -- | -- |
| BK79 to SI87 | -- | -- | 307,310 | 175 | -- | 34,112, 289,326 |
| SI87 to BE89 | -- | 163 | 45,47, 273,278 | -- | -- | 6,58, 220 |
| SI87 to BE92 | -- | -- | 47 | 96 | -- | -- |
| BE92 to WU95 | -- | 128,165 | 45,47, 278 | 96 | -- | -- |
| WU95 to SY97 | 133 | -- | 45,273 | 96,121 | 92 | 233,313 |
| SY97 to FU02 | -- | -- | 283 | 209 | 83,92 | -- |

Note： Sites reported by experimentally validated are marked with a triangle (▼). No sites identified by Koel et al. were found.

**Table S3.** Model inferred sequence position with normalized score greater than 1 in antigenic cluster transitions for A(H3N2), 2010-2022.

| Rank | Cluster 1 to 2 | | | | Cluster 2 to 3 | | | |
|---|---|---|---|---|---|---|---|---|
| | Position | Epitope | Koel site | Score | Position | Epitope | Koel site | Score |
| 1 | 48 | C | No | 4.12 | 131 | A | No | 4.63 |
| 2 | 156 | B | Yes | 4.01 | 261 | E | No | 3.82 |
| 3 | 131 | A | No | 3.75 | 48 | C | No | 3.74 |
| 4 | 142 | A | No | 3.70 | 214 | D | No | 3.35 |
| 5 | 173 | D | No | 3.40 | 183 | Others | No | 3.30 |
| 6 | 140 | A | No | 3.33 | 156 | B | Yes | 3.16 |
| 7 | 261 | E | No | 3.24 | 173 | D | No | 2.95 |
| 8 | 214 | D | No | 3.02 | 140 | A | No | 2.75 |
| 9 | 183 | Others | No | 3.01 | 172 | D | No | 2.65 |
| 10 | 158 | B | Yes | 2.77 | 142 | A | No | 2.51 |
| 11 | 157 | B | No | 2.65 | 192 | B | No | 2.49 |
| 12 | 57 | E | No | 2.54 | 158 | B | Yes | 2.24 |
| 13 | 160 | B | No | 2.51 | 122 | A | No | 2.13 |
| 14 | 172 | D | No | 2.18 | 160 | B | No | 2.03 |
| 15 | 122 | A | No | 1.77 | 212 | D | No | 1.94 |
| 16 | 212 | D | No | 1.76 | 159 | B | Yes | 1.72 |
| 17 | 92 | E | No | 1.74 | 195 | Others | No | 1.68 |
| 18 | 278 | C | No | 1.44 | 157 | B | No | 1.64 |
| 19 | 186 | B | No | 1.33 | 193 | B | Yes | 1.42 |
| 20 | 312 | C | No | 1.31 | 144 | A | No | 1.39 |
| 21 | 124 | A | No | 1.27 | 312 | C | No | 1.33 |
| 22 | 144 | A | No | 1.25 | 92 | E | No | 1.26 |
| 23 | 137 | A | No | 1.11 | 33 | Others | No | 1.16 |
| 24 | 194 | B | No | 1.00 | 189 | B | Yes | 1.11 |
| 25 | -- | -- | -- | -- | 194 | B | No | 1.09 |
| 26 | -- | -- | -- | -- | 137 | A | No | 1.04 |

**Table S4.** Model inferred sequence position with normalized score greater than 1 in antigenic cluster transitions for A(H1N1)pdm.

| Rank | Position | Epitope | Score |
|---|---|---|---|
| 1 | 236 | Ca1 | 11.93 |
| 2 | 120 | Others | 5.98 |
| 3 | 205 | Ca1 | 4.93 |
| 4 | 125 | Sa | 3.81 |
| 5 | 222 | Ca2 | 3.14 |
| 6 | 185 | Sb | 3.03 |
| 7 | 166 | Ca1 | 2.89 |
| 8 | 84 | Others | 2.82 |
| 9 | 43 | Others | 2.68 |
| 10 | 277 | Others | 2.57 |
| 11 | 164 | Sa | 2.57 |
| 12 | 282 | Others | 1.71 |
| 13 | 119 | Others | 1.67 |
| 14 | 286 | Others | 1.58 |
| 15 | 127 | Others | 1.15 |

**Table S5.** Relative risk of identifying a key site in each antigenic epitope than chance during antigenic cluster transitions.

| Cluster transition | Antigenic epitopes | Relative risk (95% CI) |
|---|---|---|
| A(H3N2) 1 to 2 | A | 5.05 (2.50, 10.21) |
| | B | 3.74 (1.71, 8.19) |
| | C | 1.52 (0.49, 4.74) |
| | D | 1.34 (0.49, 3.66) |
| | E | 1.87 (0.61, 5.73) |
| | Koel | 3.92 (1.14, 13.44) |
| A(H3N2) 2 to 3 | A | 4.00 (1.87, 8.52) |
| | B | 5.18 (2.78, 9.65) |
| | C | 0.94 (0.23, 3.74) |
| | D | 1.23 (0.45, 3.36) |
| | E | 1.15 (0.29, 4.54) |
| | Koel | 9.04 (4.98, 16.41) |
| A(H1N1)pdm 1 to 2 | Sa | 3.35 (0.85, 13.16) |
| | Sb | 1.82 (0.26, 12.65) |
| | Ca1 | 5.95 (2.01, 17.58) |
| | Ca2 | 2.72 (0.41, 18.20) |
| | Cb | Not estimated |

### Supplementary Figures

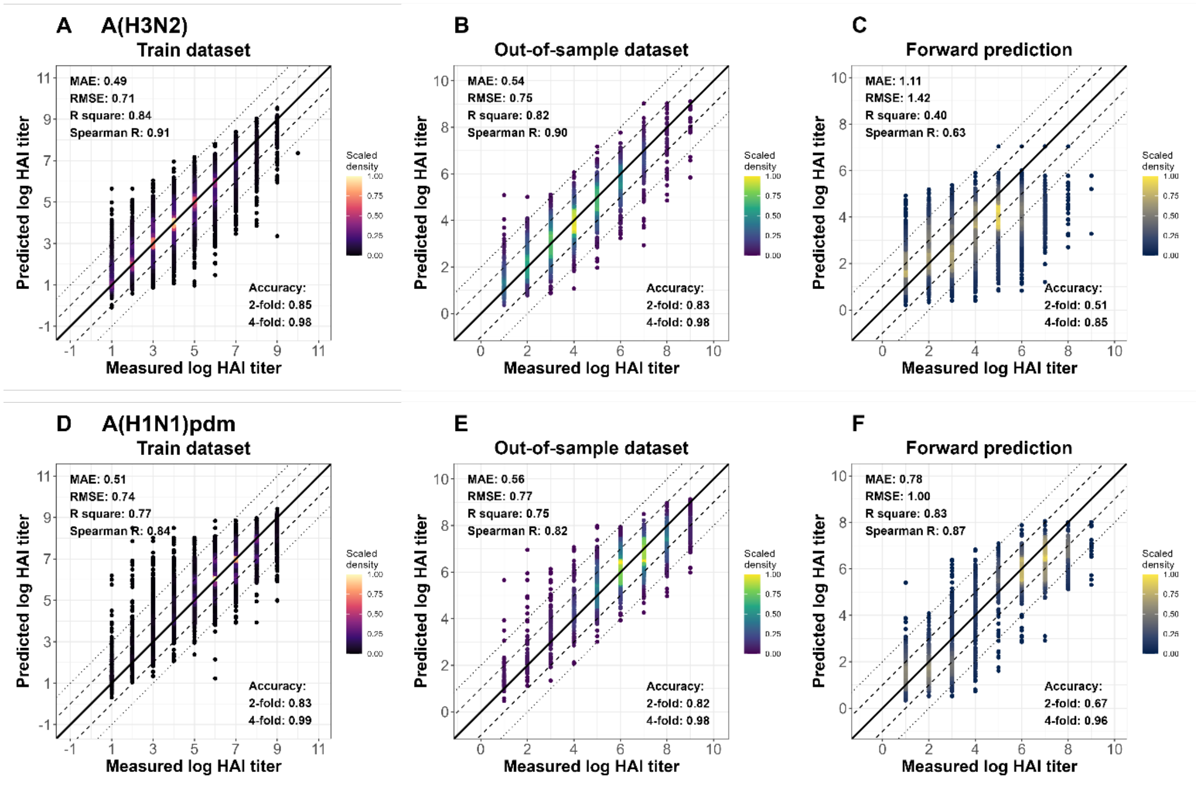


**Figure S1. Comparison of predicted and measured hemagglutination inhibition (HAI) titers for influenza A across training, ten-fold out-of-sample, and forward prediction datasets.** Colored scatter plots represent the predicted versus measured log HAI titers, with scaled density indicated by the color gradient, where higher values represent regions with more predictions falling in that location. Solid lines represent perfect predictions, dashed lines indicate errors within ±2-fold, and dotted lines mark errors within ±4-fold. Model performance is evaluated using mean absolute error (MAE), root mean square error (RMSE), $R^2$, Spearman correlation (R), and the proportion of predictions within 2-fold (1 log unit) or 4-fold (2 log units) of the measured values. (**A**) A(H3N2) HAI titer predictions for the train dataset (2012–2020), which includes 42,893 titrations (2,915 and 89 identical test and reference viruses). (**B**) A(H3N2) HAI titer predictions for the out-of-sample dataset (2012–2020), comprising 4,766 titrations (1,850 and 78 identical test and reference viruses) that are not included in the training dataset. (**C**) A(H3N2) HAI titer predictions for the forward prediction dataset (2021–2022), containing 16,642 titrations (1,250 and 32 identical test and reference viruses). (**D**) A(H1N1)pdm HAI titer predictions for the train dataset (2012–2020), including 54,712 titrations (4,490 and 38 identical test and reference viruses). (**E**) A(H1N1)pdm HAI titer predictions for the out-of-sample dataset (2012–2020), consisting of 6,079 titrations (2,812 and 38 identical test and reference viruses) that are not included in the training dataset. (**F**) A(H1N1)pdm HAI titer predictions for the forward prediction dataset (2021–2022), comprising 4,714 titrations (324 and 21 identical test and reference viruses).

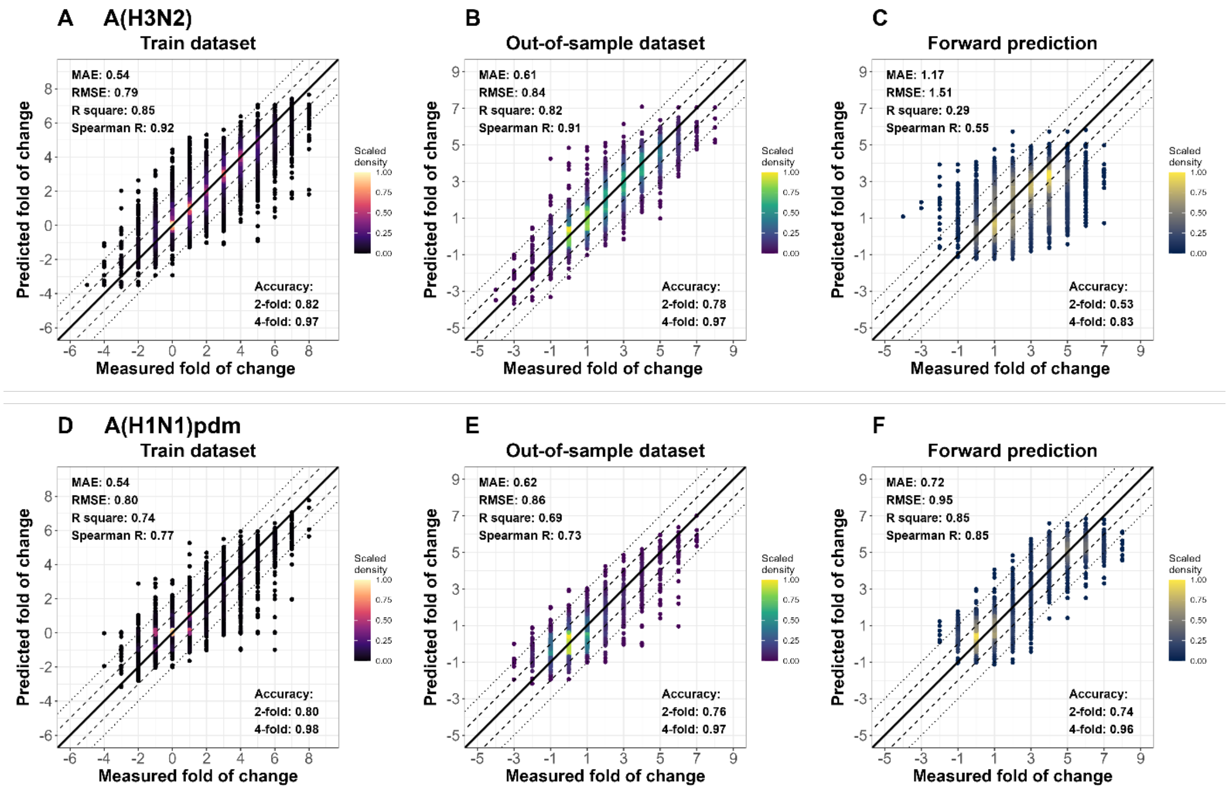


**Figure S2. Comparison of predicted and measured fold-of-change in hemagglutination inhibition (HAI) titers for influenza A across training, ten-fold out-of-sample, and forward prediction datasets.** Colored scatter plots represent the predicted versus measured fold change of HAI titers, with scaled density indicated by the color gradient, where higher values represent regions with more predictions falling in that location. Solid lines represent perfect predictions, dashed lines indicate errors within ±2-fold, and dotted lines mark errors within ±4-fold. Model performance is evaluated using mean absolute error (MAE), root mean square error (RMSE), $R^2$, Spearman correlation (R), and the proportion of predictions within 2-fold (1 log unit) or 4-fold (2 log units) of the measured values. (**A**) A(H3N2) fold-change predictions for the train dataset (2012–2020), which includes 42,893 titrations (2,915 and 89 identical test and reference viruses). (**B**) A(H3N2) fold-change predictions for the out-of-sample dataset (2012–2020), comprising 4,766 titrations (1,850 and 78 identical test and reference viruses) that are not included in the training dataset. (**C**) A(H3N2) fold-change predictions for the forward prediction dataset (2021–2022), containing 16,642 titrations (1,250 and 32 identical test and reference viruses). (**D**) A(H1N1)pdm fold-change predictions for the train dataset (2012–2020), including 54,712 titrations (4,490 and 38 identical test and reference viruses). (**E**) A(H1N1)pdm fold-change predictions for the out-of-sample dataset (2012–2020), consisting of 6,079 titrations (2,812 and 38 identical test and reference viruses) that are not included in the training dataset. (**F**) A(H1N1)pdm fold-change predictions for the forward prediction dataset (2021–2022), comprising 4,714 titrations (324 and 21 identical test and reference viruses).

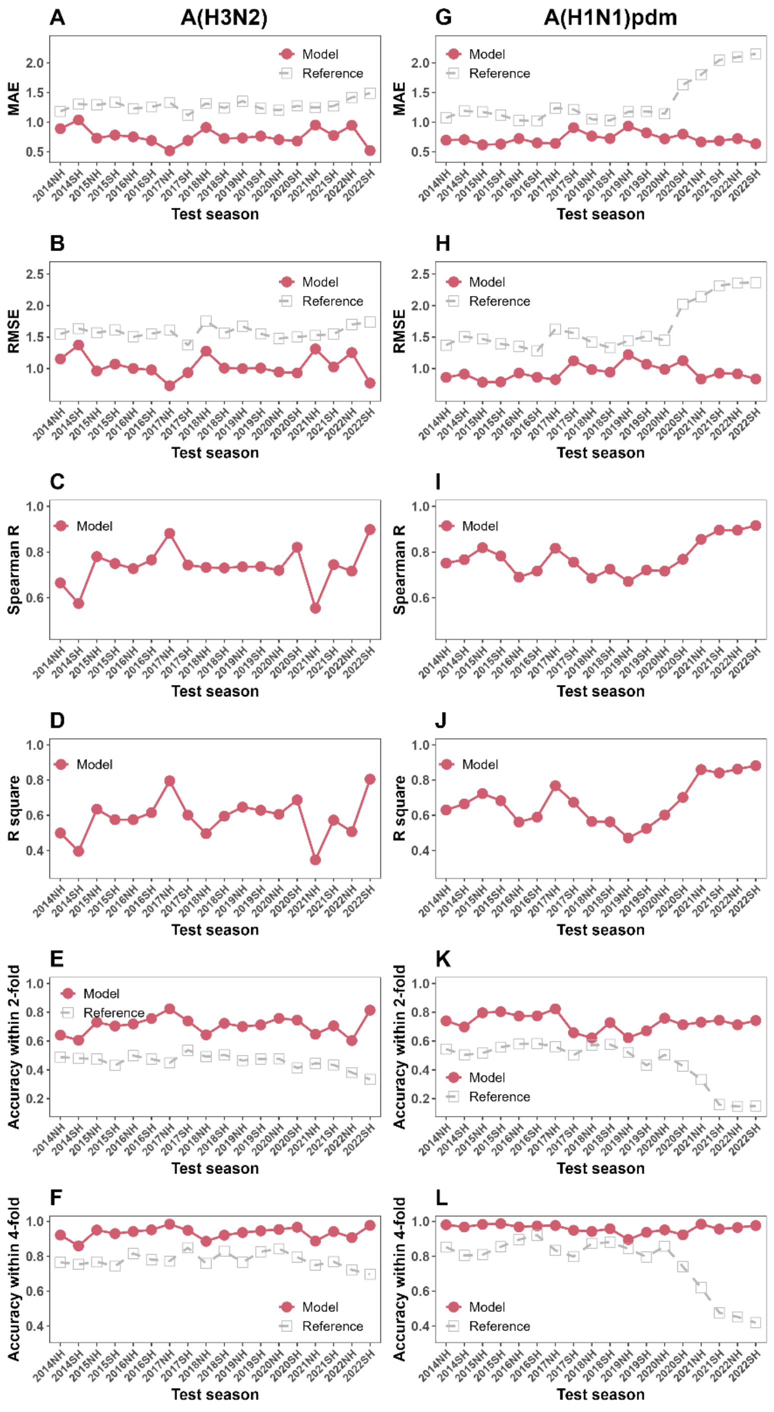


**Figure S3. Season-by-season model performance on influenza A hemagglutination inhibition (HAI) titer predictions.** Model performance was evaluated using various statistical metrics on a rolling test dataset applied season-by-season. Panels **A–F** show A(H3N2) results; **G–L** show A(H1N1)pdm. Metrics (from top to bottom) are mean absolute error, root-mean-square error, Spearman r, $R^2$, and the proportions of predictions within 2- and 4-fold of observed titers.

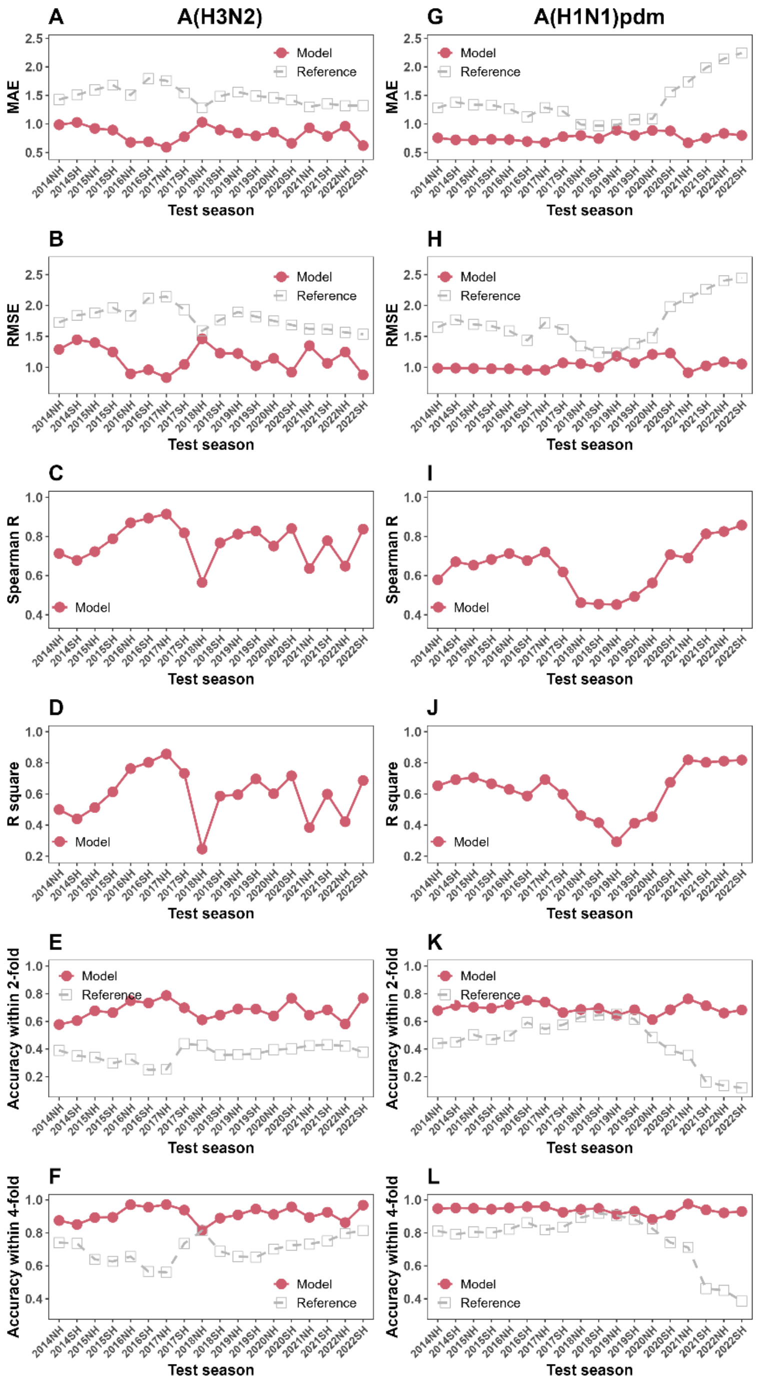


**Figure S4. Season-by-season model performance on influenza A hemagglutination inhibition (HAI) fold-of-change predictions.** Model performance was evaluated using various statistical metrics on a rolling test dataset applied season-by-season. Panels **A–F** show A(H3N2) results; **G–L** show A(H1N1)pdm. Metrics (from top to bottom) are mean absolute error, root-mean-square error, Spearman r, $R^2$, and the proportions of predictions within 2- and 4-fold of observed titers.

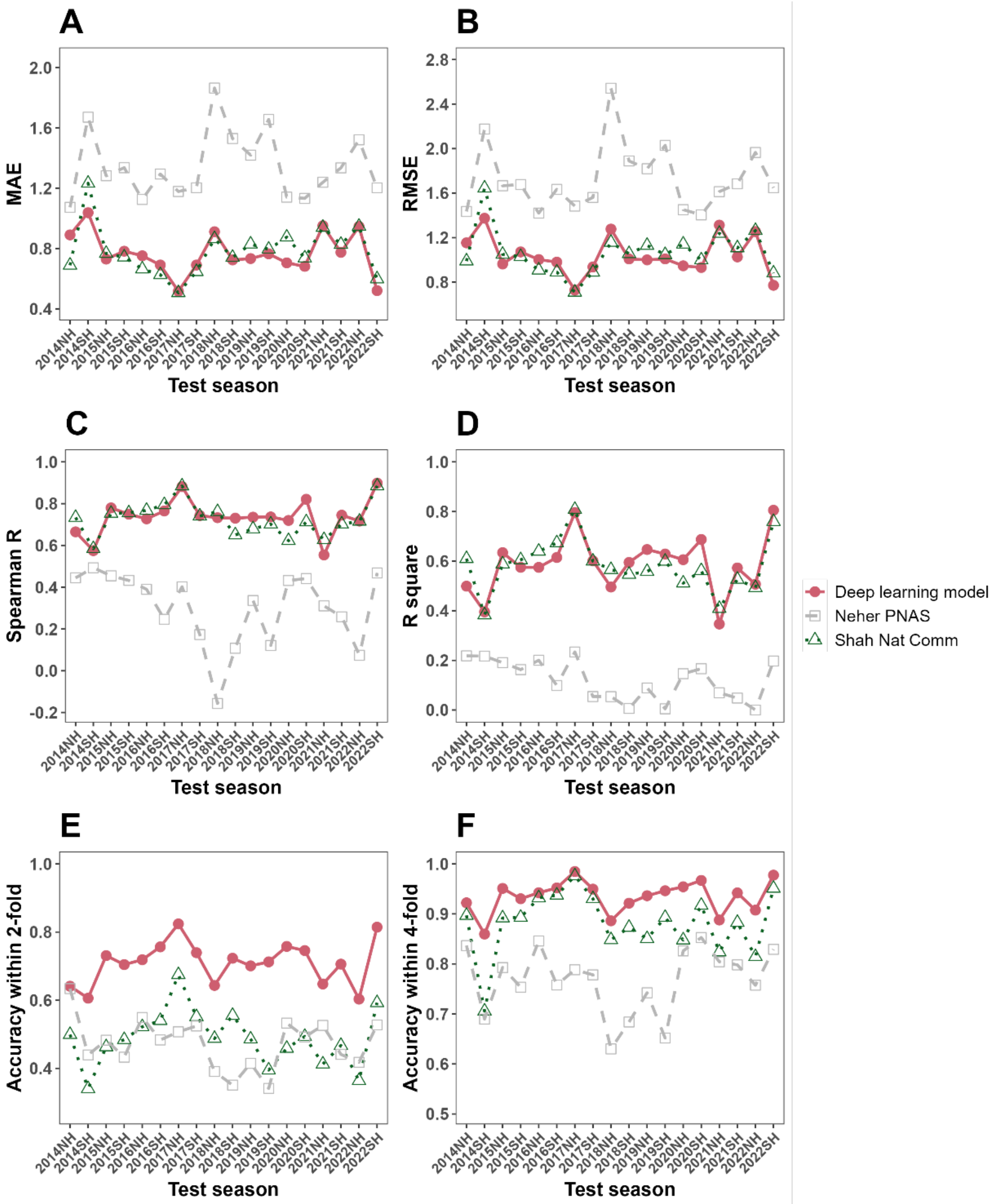


**Figure S5. Season-by-season model performance of influenza A(H3N2) hemagglutination inhibition (HAI) titer predictions, compared to existing models.** Statistical metrics were compared among our deep learning model, Neher's substitution model and Shah's Adaboost model. Model performances were evaluated on a season-by-season rolling test. Logarithm transformed HAI titer was assessed, with the geometric mean titer of the examined report being used as reference. (**A**) Mean absolute error (MAE) for A(H3N2) predictions. (**B**) Root means square error (RMSE) for A(H3N2) predictions. (**C**) Spearman correlation (R) for A(H3N2) predictions. (**D**) $R^2$ values for A(H3N2) predictions. (**E**) Accuracy proportion of predictions within 2-fold for A(H3N2) predictions. (**F**) Accuracy proportion of predictions within 4-fold for A(H3N2) predictions.

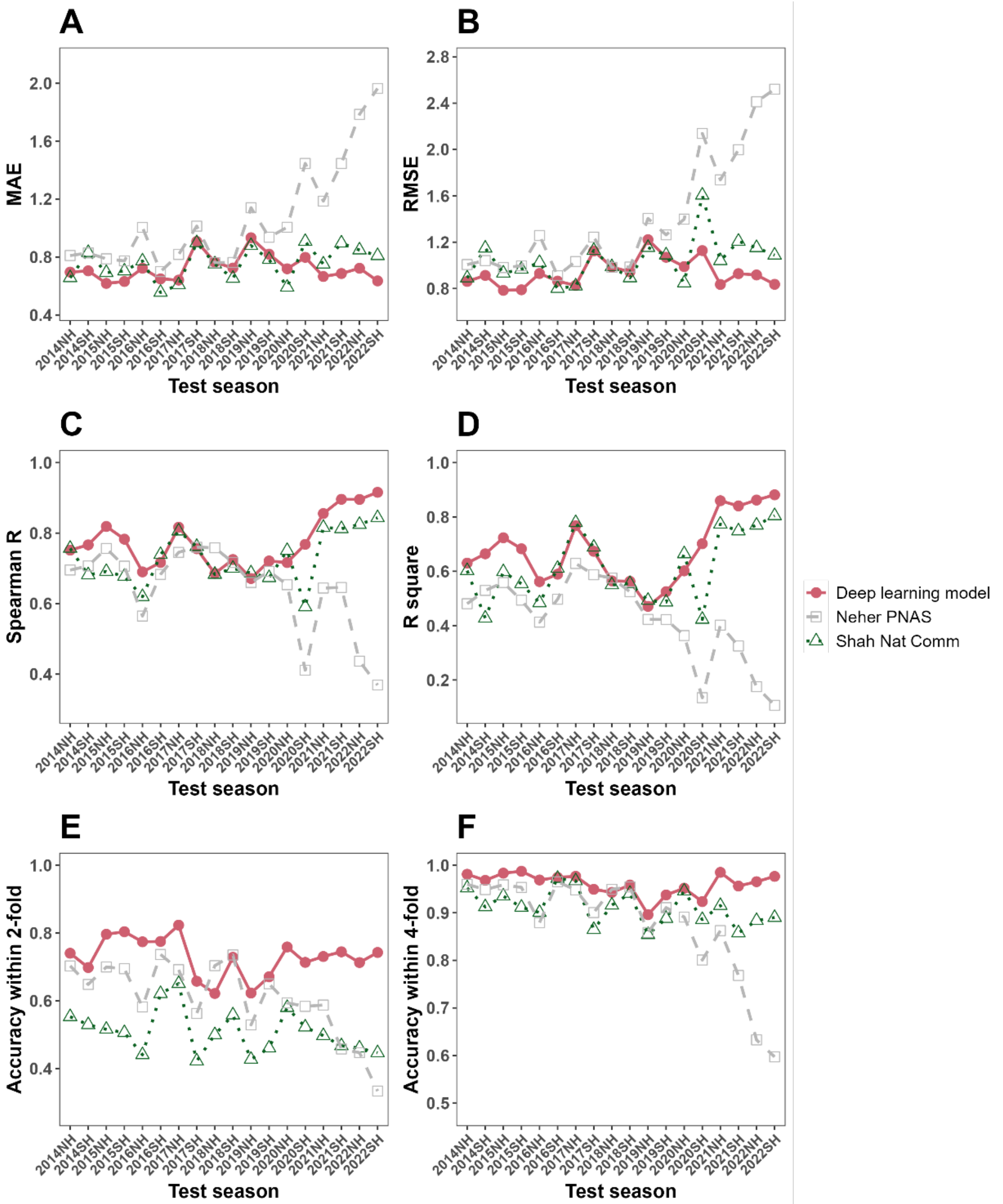


**Figure S6. Season-by-season model performance of influenza A(H1N1)pdm hemagglutination inhibition (HAI) titer predictions, compared to existing models.** Statistical metrics were compared among our deep learning model, Neher's substitution model and Shah's Adaboost model. Model performances were evaluated on a season-by-season rolling test. Logarithm transformed HAI titer was assessed, with the geometric mean titer of the examined report being used as reference. (**A**) Mean absolute error (MAE) for A(H1N1)pdm predictions. (**B**) Root means square error (RMSE) for A(H1N1)pdm predictions. (**C**) Spearman correlation (R) for A(H1N1)pdm predictions. (**D**) $R^2$ values for A(H1N1)pdm predictions. (**E**) Accuracy proportion of predictions within 2-fold for A(H1N1)pdm predictions. (**F**) Accuracy proportion of predictions within 4-fold for A(H1N1)pdm predictions.

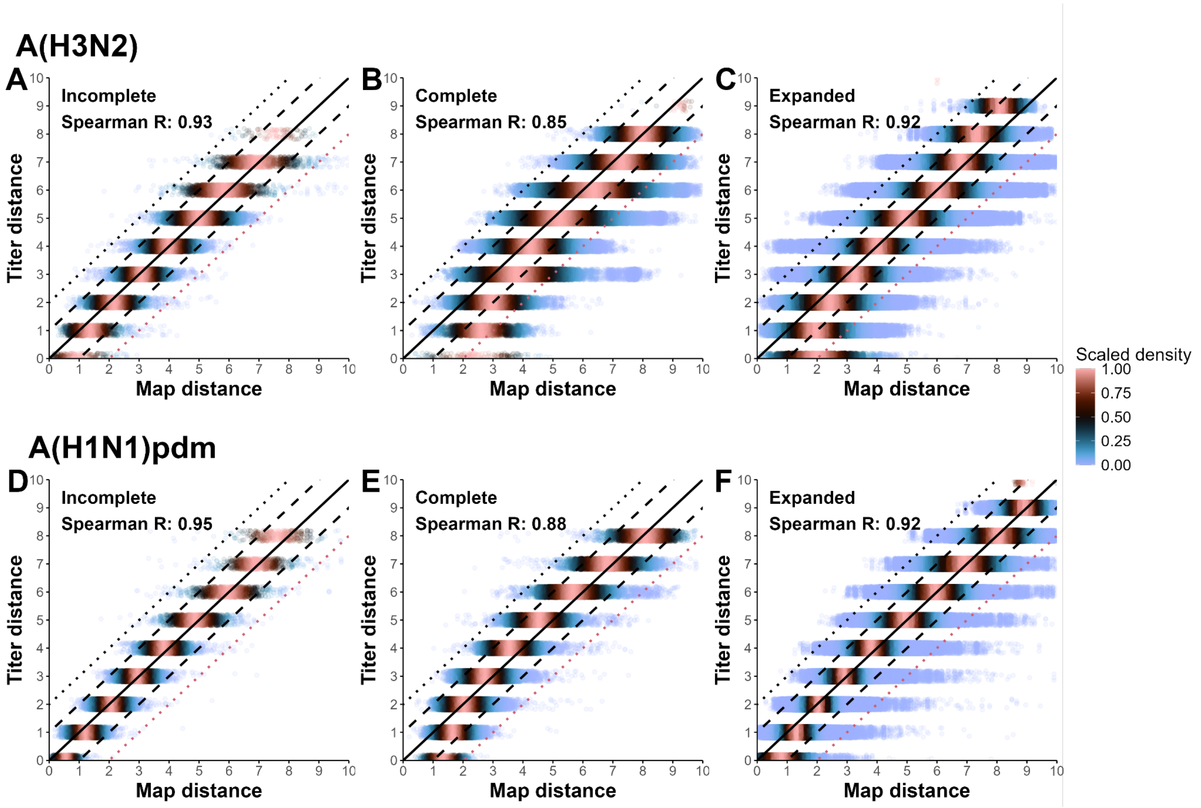


**Figure S7. Comparison of targeted and antigenic distances between antisera and viruses.** Titer distance represents the log-difference between column maximum titers and measured HAI titers for each antiserum. Antigenic distance is the Euclidean distance calculated from the map coordinates of the virus and antiserum. Solid, dashed, and dotted lines indicate errors of 0, ±1, and ±2 log units, respectively. Antigenic maps were estimated using incomplete measured data only (**A**, **D**), complete predicted and measured data (**B**, **E**) or expanded predicted data (**C**, **F**). Panels **A-C** show results for A(H3N2), and panels **D-F** for A(H1N1)pdm.

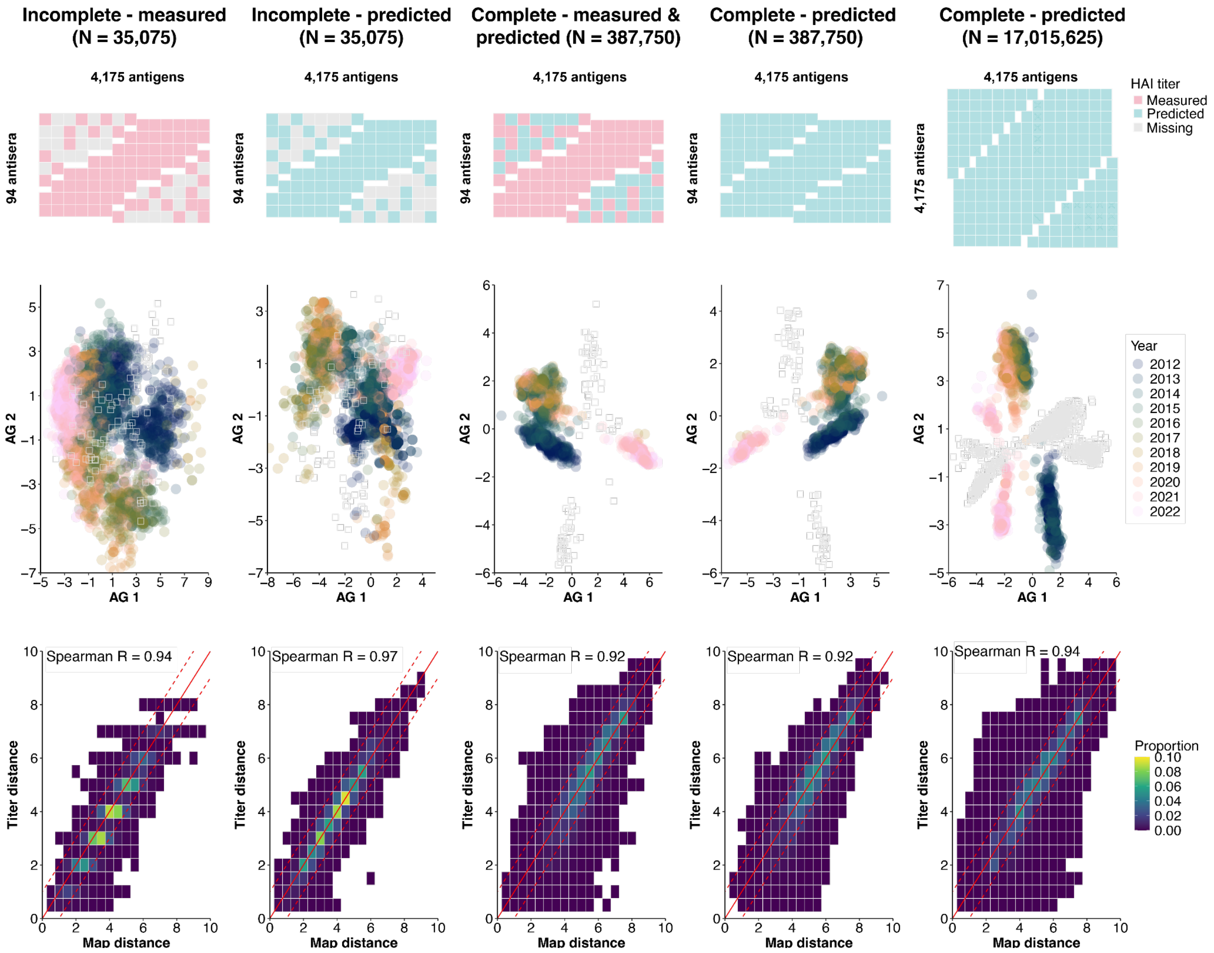


**Figure S8. Comparison of incomplete, complete, and expanded antigenic maps for A(H3N2).** The top row shows conceptual illustrations of HAI titer coverage for 94 reference viruses (rows) and 4,175 test viruses (columns) under different titer matrices. From left to right, the columns represent: (1) incomplete measured data, (2) incomplete predicted data, (3) complete measured and predicted data, (4) complete predicted data, and (5) expanded predicted data. The middle row shows two-dimensional antigenic maps generated by MDS, with each test virus colored by isolation year (2012–2022). The bottom row compares pair-wise titer distances with map distances. The solid red line indicates identical distance estimates, while the dashed red lines represent a one-log-unit difference between the two maps. The Spearman correlation coefficient (r) is reported in each panel.

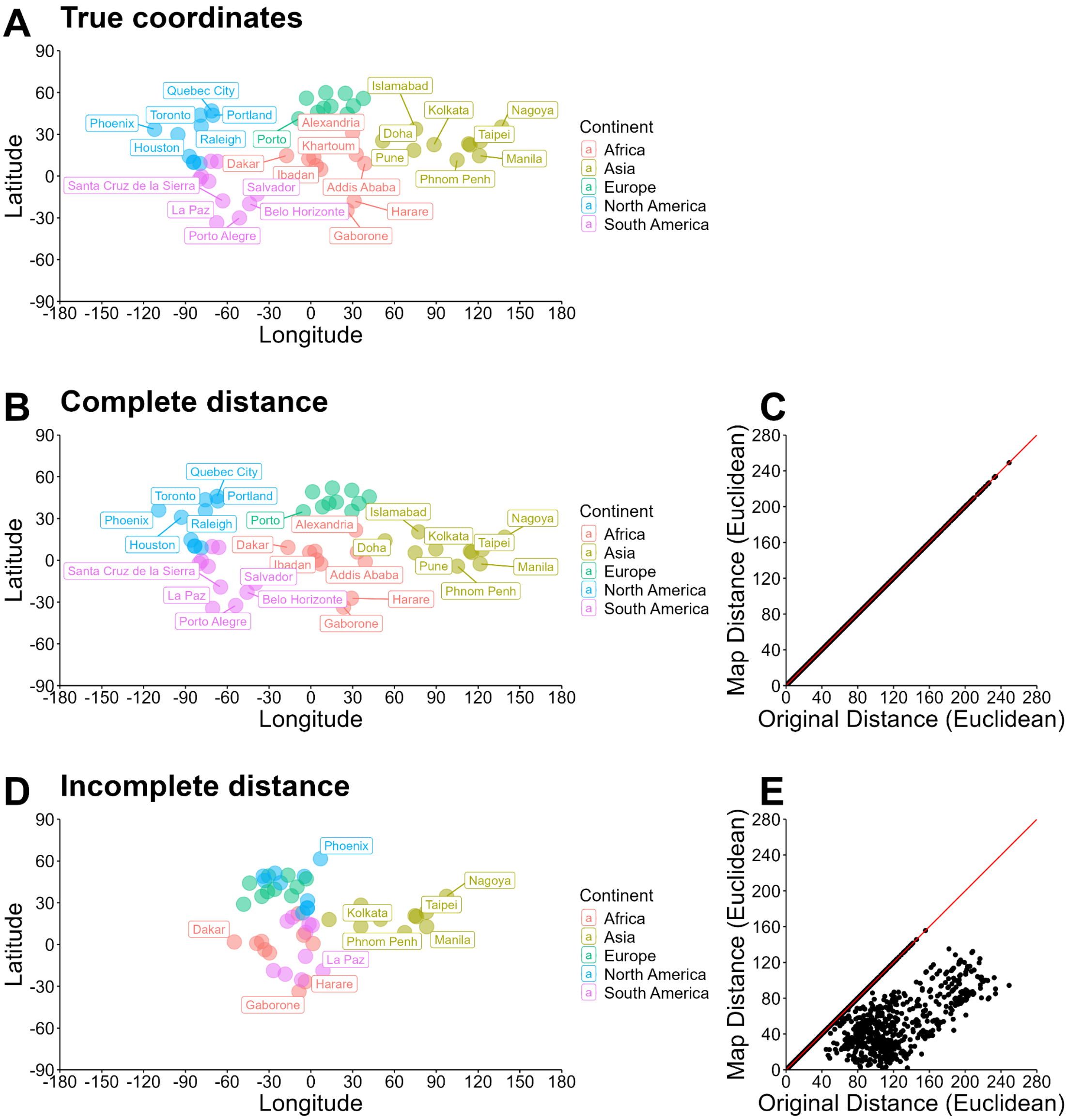


**Figure S9. Geographic versus MDS positions for 50 world cities reconstructed from incomplete and complete distance matrices.** (**A**) True geographic coordinates of 50 randomly selected cities, with 10 for each of Africa, Asia, Europe, North America and South America and color-coded by continents. (**B**) Two-dimensional MDS map reconstructed from the complete matrix of pair-wise Euclidean distance distances. (**C**) Scatterplot of MDS-inferred distances versus true geographic distances for the complete distance matrix. The red line indicates perfect agreement. (**D**) Two-dimensional MDS map reconstructed from the incomplete matrix retaining only the shortest 20 % of pair-wise distances. (**E**) Scatterplot of MDS-inferred distances versus true geographic distances for the incomplete matrix of pair-wise distances.

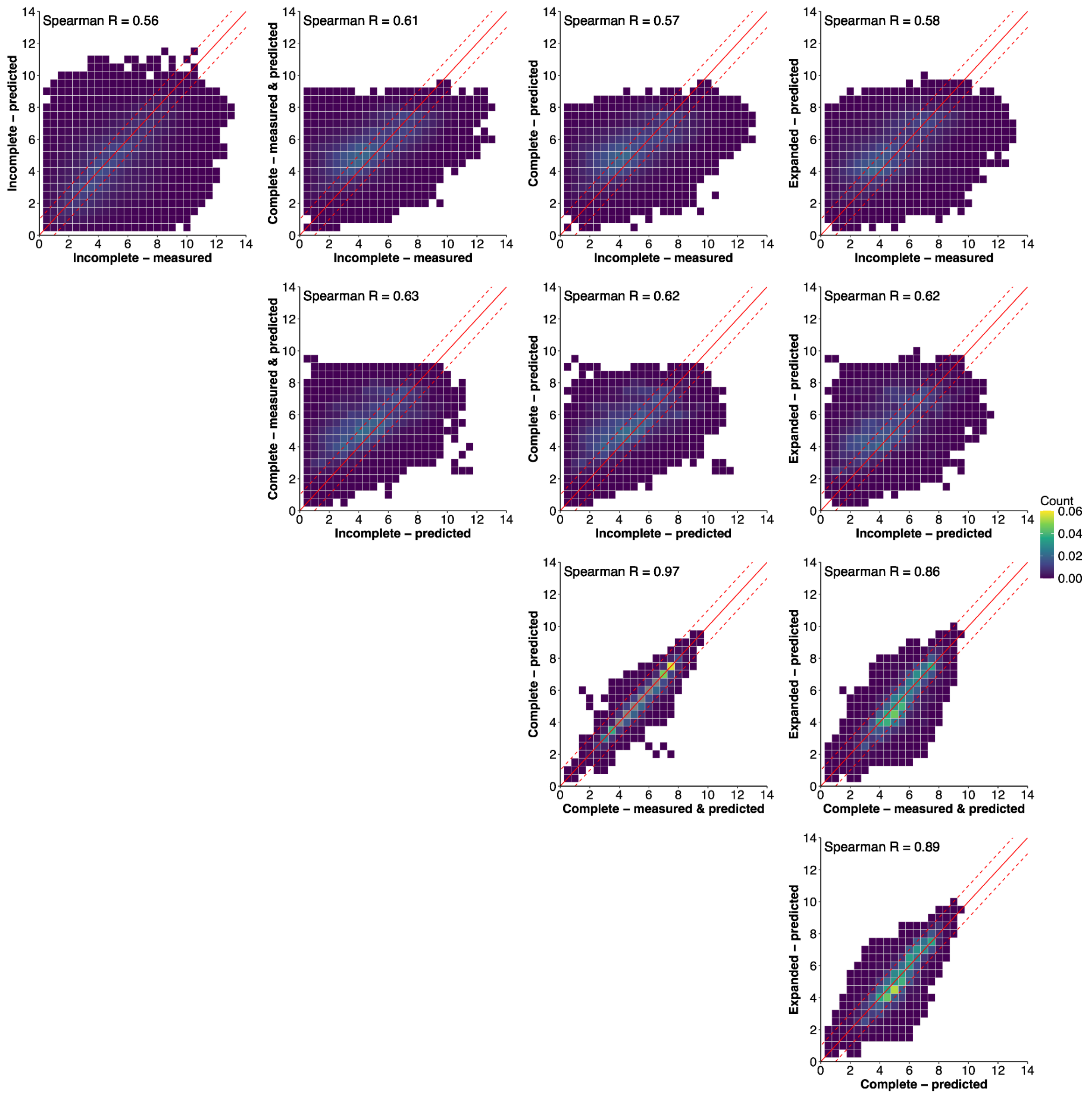


**Figure S10. Comparison of predicted antigenic distances for A(H3N2) under incomplete, complete, and expanded titer matrices.** Each panel shows pair-wise distance comparisons from two mapping approaches. Five datasets were used: (1) incomplete measured titers, (2) incomplete predicted titers, (3) a complete matrix of measured titers with missing values filled by predicted titers, (4) a complete matrix of predicted titers, and (5) an expanded matrix of predicted titers. The solid red line indicates identical distance estimates, while the dashed red lines represent a one-log-unit difference between the two maps. The Spearman correlation coefficient (r) is reported in each panel.

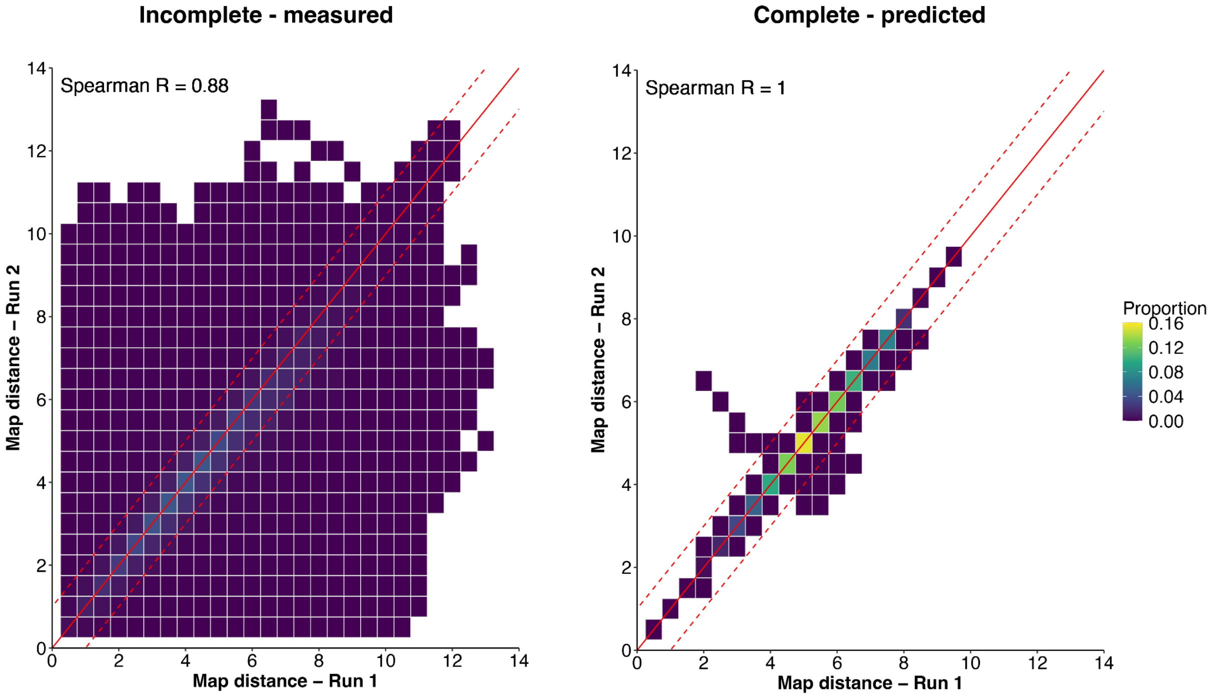


**Figure S11. Comparison of predicted antigenic distances for A(H3N2) using incomplete and complete titer matrices across independent runs.** Complete matrix contains HAI titers predicted from the deep learning model. The solid red line marks perfect agreement between distance estimates, and the dashed red lines indicate a ±1 log-unit difference. The Spearman correlation coefficient (r) is shown in each panel.

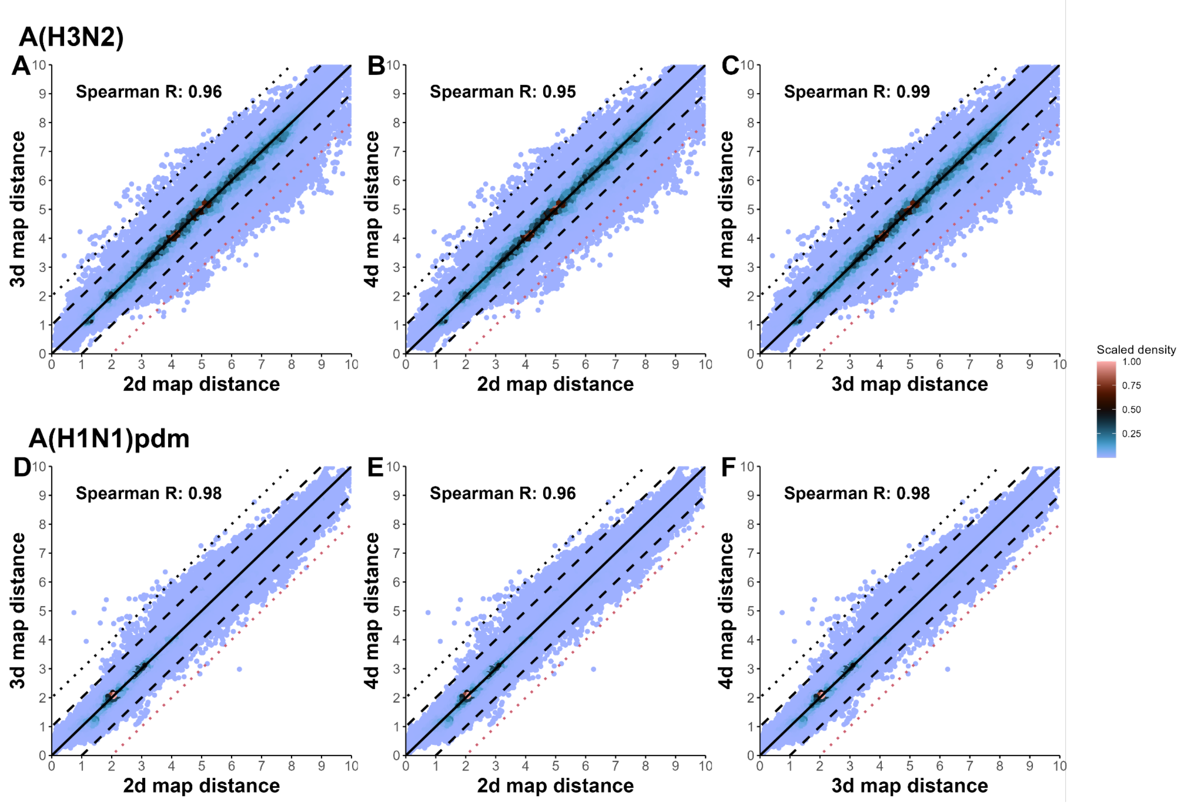


**Figure S12. Comparison of antigenic distances among different dimensionalities.** We used expanded titer matrices to create antigenic maps in two, three, and four dimensions. Higher values in the scaled density represent regions with more predictions falling in that location (**A**-**C**) Comparison of map distances for A(H3N2). (**D**-**F**) Comparison of map distance for A(H1N1)pdm.

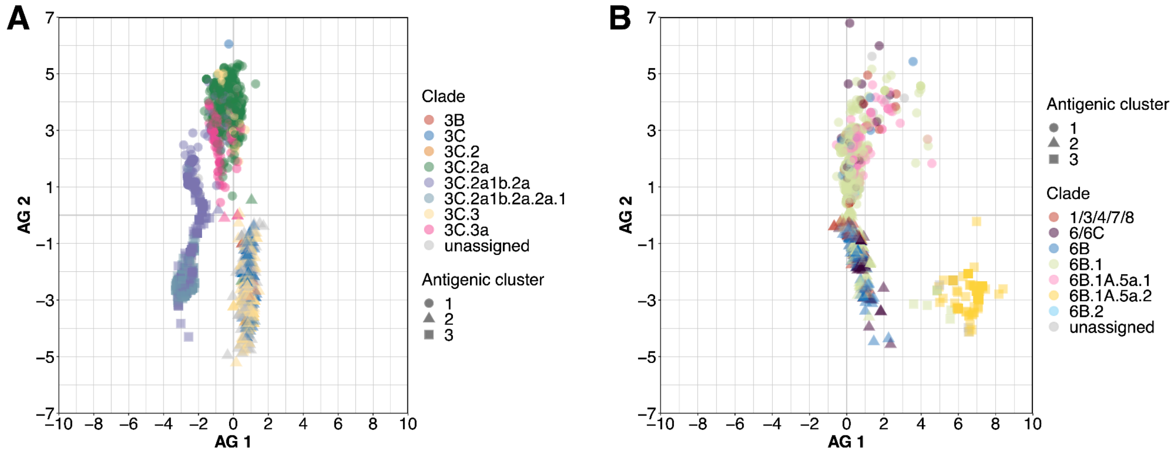


**Figure S13. Antigenic maps colored by genetic clades of viruses.** The map estimates are the same in Figure 4 A, D. **(A)** A(H3N2). **(B)** A(H1N1)pdm.

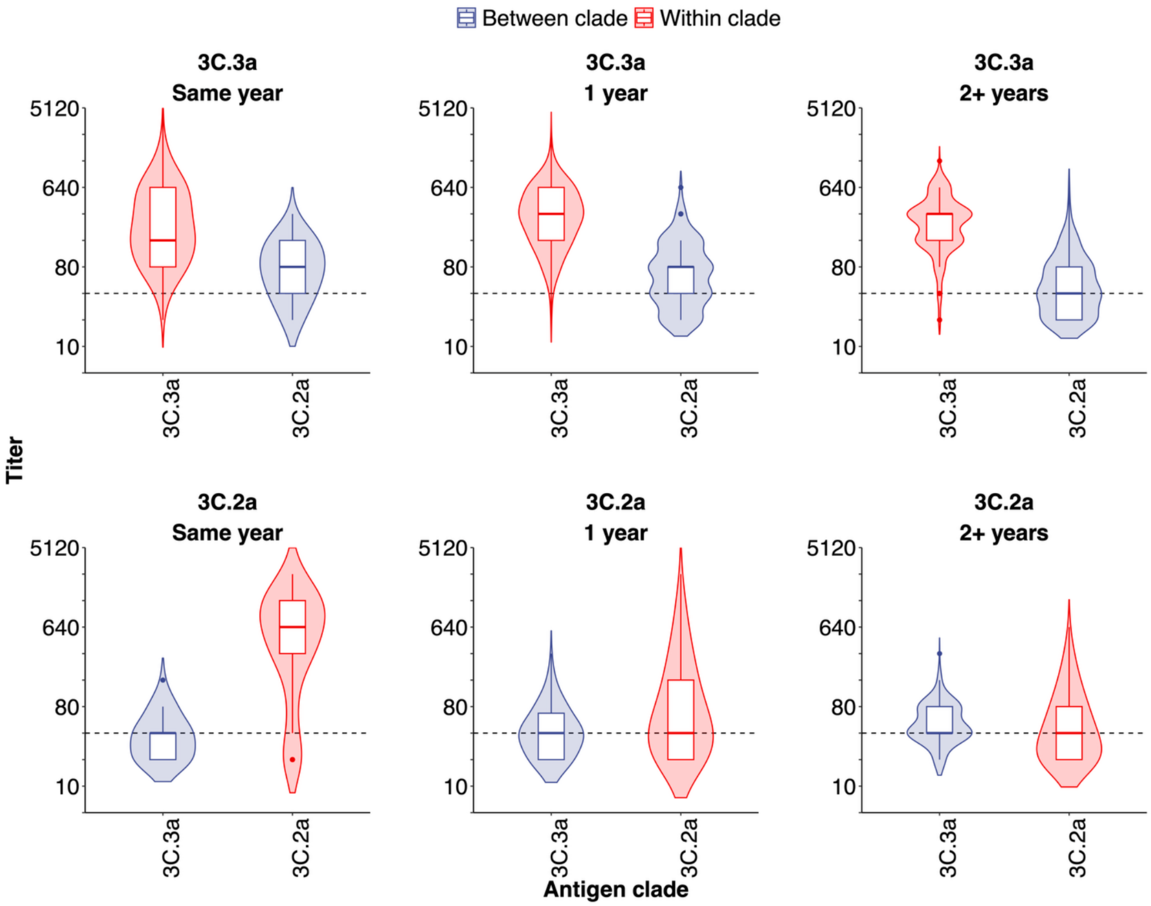


**Figure S14. Cross-reactivity between A(H3N2) 3C.2a and 3C.3a subclades**. Only viruses from the two subclades were included. Titers were stratified by the difference in their isolation years. The x-axis shows the subclade of the test viruses, and each panel corresponds to the subclade of the antiserum. Dashed horizontal lines indicate a titer for 40.

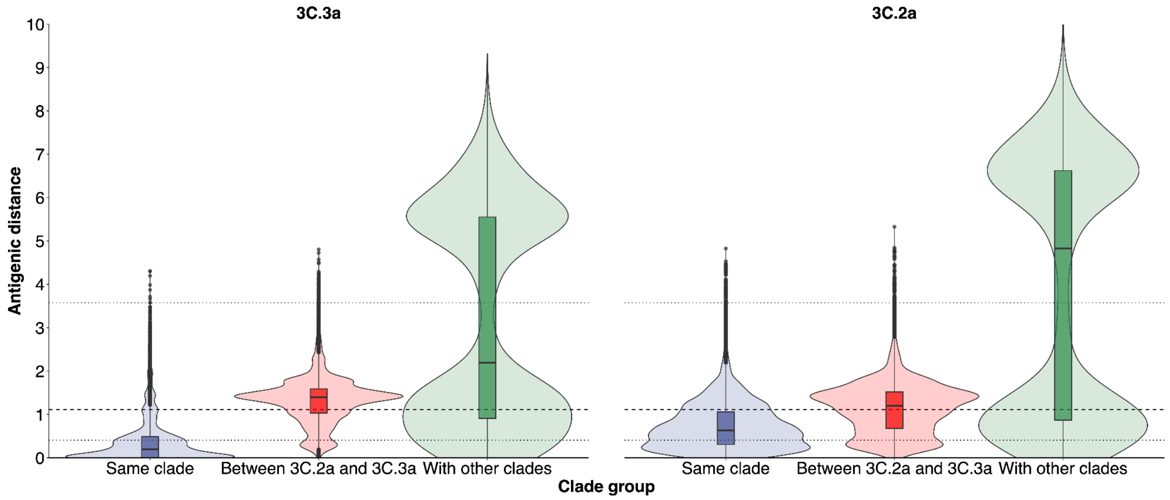


**Figure S15. Antigenic distance between A(H3N2) 3C.2a and 3C.3a subclades.** All available viruses from the expanded maps (Figure 4A) are shown, with antigenic distances stratified by the subclade of test viruses (x-axis). Each panel corresponds to the subclade of viruses used to raise antiserum. Dashed and dotted horizontal lines denote the median and interquartile range of antigenic distances between viruses isolated one year apart.

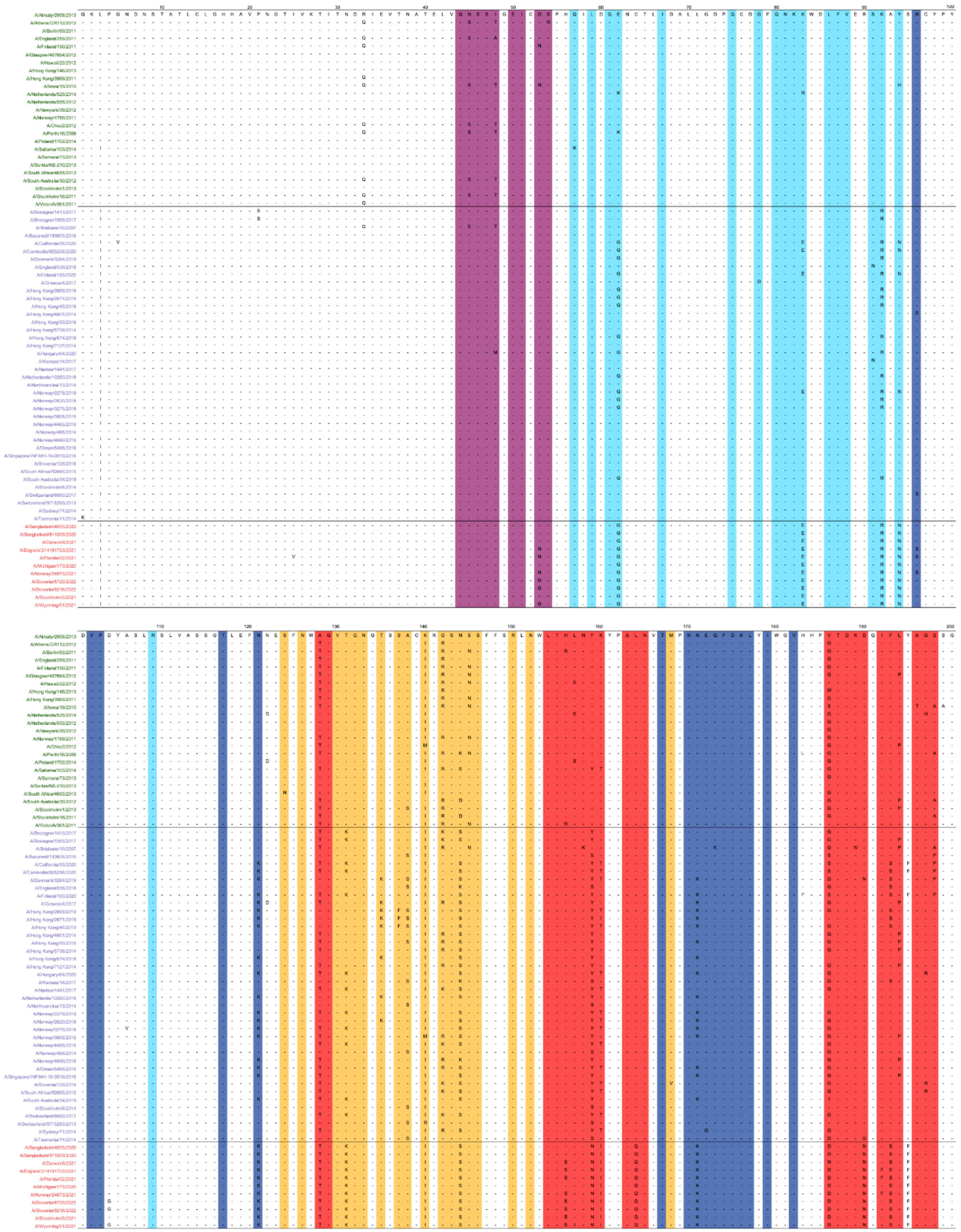

**Figure S16. Multiple sequence alignment for reference A(H3N2) viruses used to assess antigenic cluster transitions.** HA1 amino acid Sequences were derived from the biannual WIC reports.

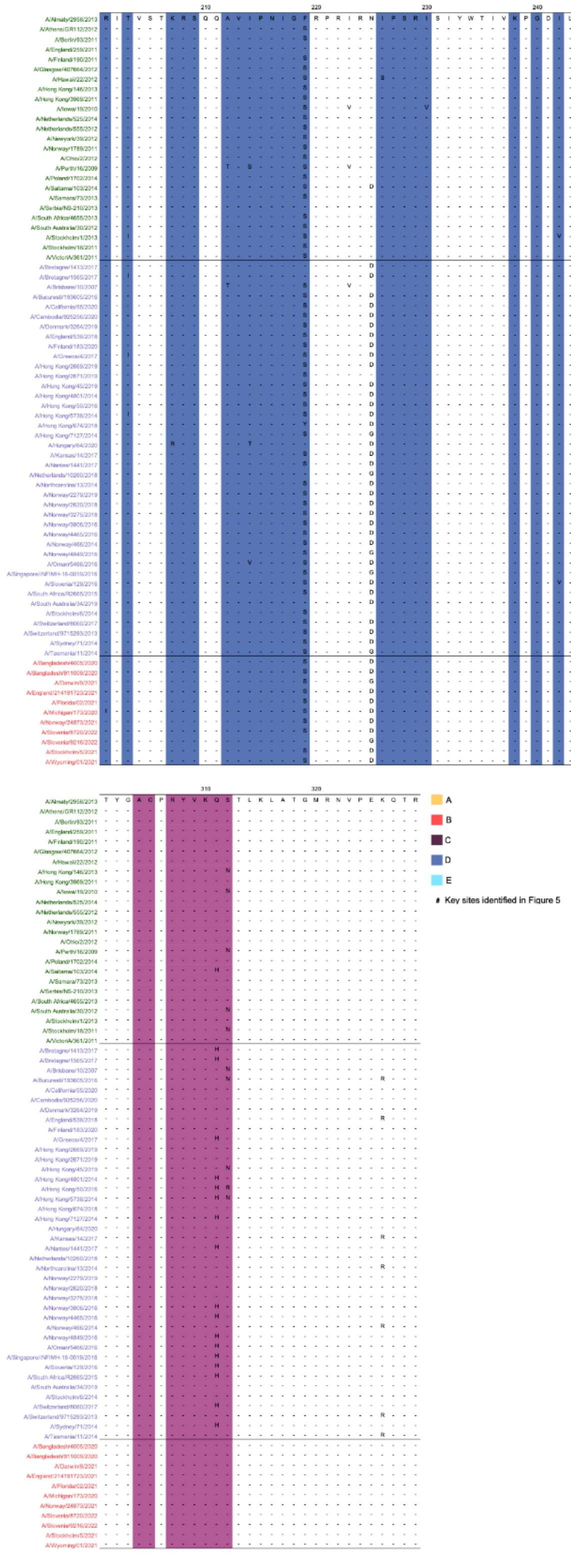


**Figure S16 (Continued). Multiple sequence alignment for reference A(H3N2) viruses used to assess antigenic cluster transitions.** HA1 amino acid Sequences were derived from the biannual WIC reports.

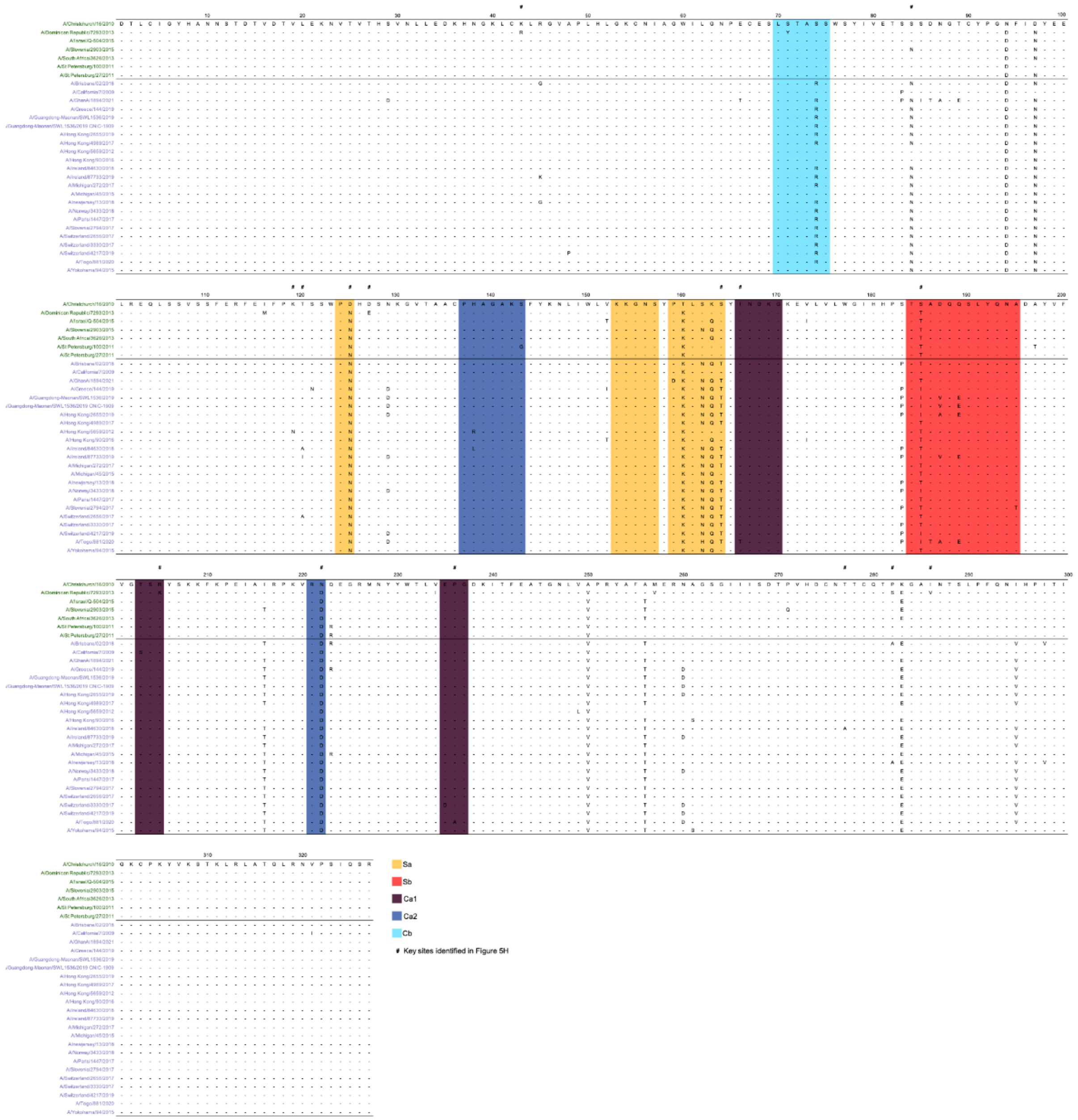


**Figure S17 Multiple sequence alignment for reference A(H1N1)pdm viruses used to assess antigenic cluster transitions.** HA1 amino acid Sequences were derived from the biannual WIC reports.

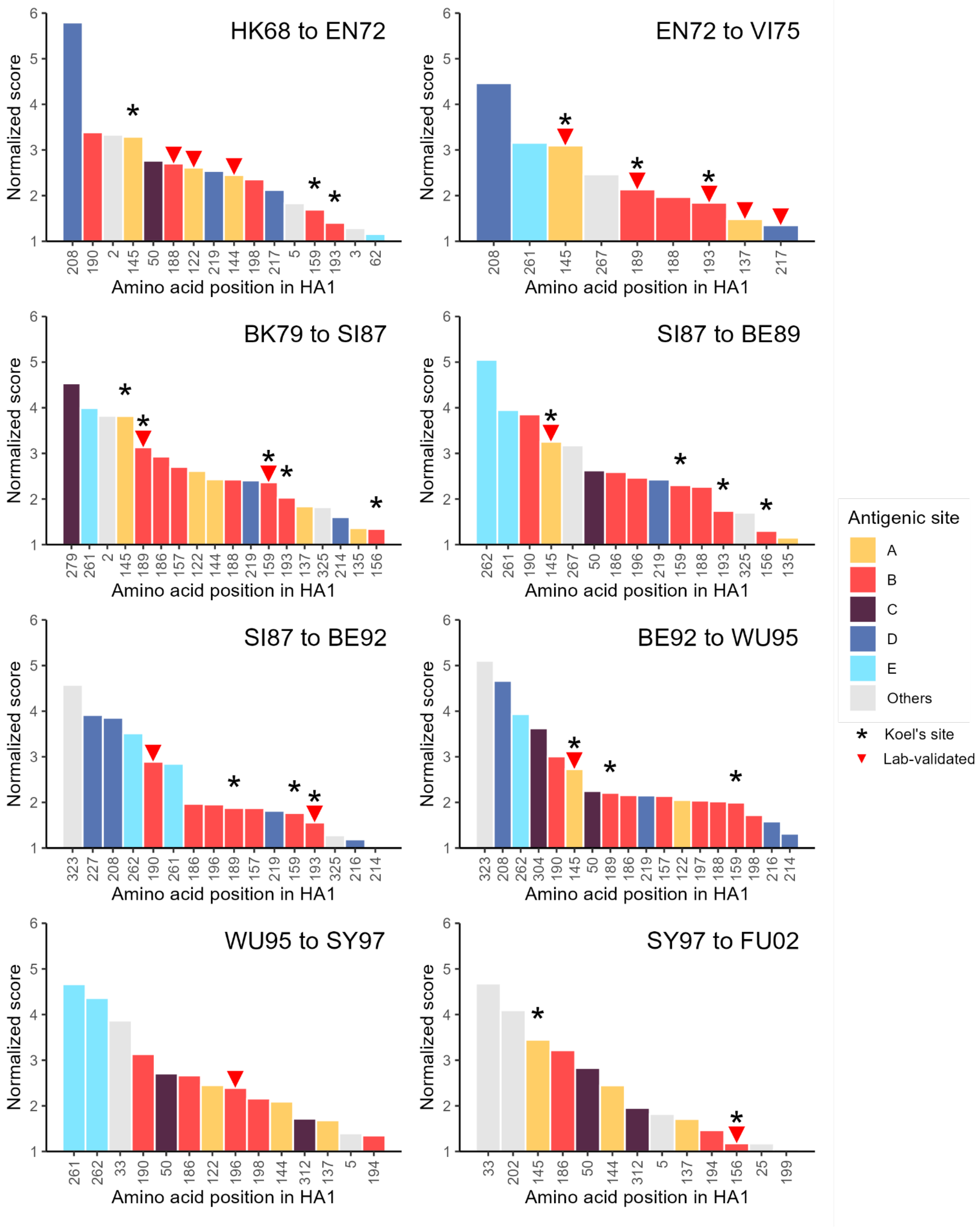


**Figure S18. Key antigenic mutation sites associated with historical A(H3N2) antigenic cluster transitions (1968–2002).** We trained a transformer model on HAI data from Smith et al. (2004) and used Grad-CAM to pinpoint mutations associated with each adjacent antigenic cluster transition. Identified sites are compared to those reported by Koel et al. (marked with a star) and to other experimentally validated mutations (red triangles). Cluster transitions were evaluated sequentially between adjacent pairs, except for Texas 1977 due to few reference viruses. Abbreviations follow the original study: HK (Hong Kong), EN (England), VI (Victoria), BK (Bangkok), SI (Sichuan), BE (Beijing), WU (Wuhan), SY (Sydney), and FU (Fujian). Two-digit numbers indicate the isolation year during the 1990s.
